# Unravelling drivers of local adaptation through Evolutionary Functional-Structural Plant modelling

**DOI:** 10.1101/2022.09.02.506361

**Authors:** Jorad de Vries, Simone Fior, Aksel Pålsson, Alex Widmer, Jake M. Alexander

## Abstract

1. Local adaptation to contrasting environmental conditions along environmental gradients is a widespread phenomenon in plant populations, yet we lack a mechanistic understanding of how individual agents of selection contribute to local adaptation.
2. Here, we developed a novel evolutionary functional-structural plant (E-FSP) model that simulates local adaptation of virtual plants along an environmental gradient. First, we validate the model by testing if it can recreate two elevational ecotypes of *Dianthus carthusianorum* occurring in the Swiss Alps. Second, we use the E-FSP model to disentangle the relative contribution of abiotic (temperature) and biotic (competition and pollination) selection pressures to elevational adaptation in *D. carthusianorum*.
3. The model reproduced the qualitative differences between the elevational ecotypes in two phenological (germination and flowering time) and one morphological trait (stalk height), as well as qualitative differences in four performance variables that emerge from GxE interactions (flowering time, number of stalks, rosette area and seed production). Our results suggest that elevational adaptation in *D. carthusianorum* is predominantly driven by the abiotic environment.
4. Our approach shows how E-FSP models incorporating physiological, ecological and evolutionary mechanisms can be used in combination with experiments to examine hypotheses about patterns of adaptation observed in the field.

## Introduction

Local adaptation to contrasting environmental conditions is a widespread phenomenon in plant populations (Leimu & Fischer, 2008), resulting from divergent selection pressures imposed by variation in environmental conditions on populations occurring across a species’ range. The outcome of local adaptation can be documented in field experiments that assess the performance of alternative ecotypes in contrasting environments, where local ecotypes are expected to outperform foreign ecotypes (Kawecki & Ebert, 2004). A wealth of experimental work has shown the pervasiveness of local adaptation (Leimu & Fischer, 2008), yet we often lack a mechanistic understanding of how individual agents of selection contribute to local adaptation along environmental gradients (Wadgymar *et al*., 2017). This is caused by individual drivers of selection acting on multiple plant traits, either directly or indirectly, and by individual traits being affected by multiple drivers of selection. The interactions between drivers of selection, such as between abiotic and biotic factors (Briscoe Runquist *et al*., 2020; Hargreaves *et al*., 2020; Paquette & Hargreaves, 2021), further complicates disentangling the role of any individual driver in shaping local adaptation.

To address this, experimental studies may be complemented by mechanistic modelling approaches (Connolly *et al*., 2017) that incorporate the eco-physiological and eco-evolutionary processes driving local adaptation. Such mechanistic modelling approaches are more commonly used in crop breeding and have proven to be powerful tools to explore the mechanistic basis of plant-environment interactions (Hammer *et al*., 2006). However, the potential for these models to simulate the ecological complexity that drives local adaptation in natural plant communities currently remains unexplored.

Function-structural plant (FSP) modelling is such a mechanistic modelling approach that integrates an explicit representation of plant structure in a 3D environment, combined with functional plant responses to that environment (Evers *et al*., 2018). The approach is particularly suited to the simulation of plant-plant interactions as it explicitly simulates the spatial heterogeneity that is inherent to species mixtures and drives competitive interactions between plants (Evers *et al*., 2019; Bongers, 2020). This explicit representation of plant form and function makes FPS modelling an excellent tool to test hypotheses about the adaptive value of functional traits in a dynamic ecological context (Bongers *et al*., 2019; de Vries *et al*., 2019; Douma *et al*., 2019).

A novel and largely unexplored application of FSP modelling is in combination with a mechanistic model of natural selection (de Vries, 2021). Such evolutionary-FSP (E-FSP) models can simulate the combined selection pressure imposed by multiple selective agents on a population of individually distinct plants that interact with each other and with the environment (de Vries, 2021). This individual-based perspective is of particular importance to mechanistically simulate natural selection, as key mechanisms that drive selection (e.g. competition for resources) are not only driven by absolute trait values, but also by trait values relative to those of neighbouring plants (Falster & Westoby, 2003; McNickle & Dybzinski, 2013). This is exemplified by competition for light, which is a pre-emptive resource (i.e. light interception by one plant also prevents light interception by other plants) whose acquisition is dependent on the height of a plant relative to the height of the surrounding vegetation, leading to competitive asymmetry (Weiner, 1990). Despite the potential for E-FSP models to simulate the mechanisms that drive local adaptation, the complexity of these mechanisms makes validation of such a model particularly challenging. As such, all E-FSP models published to date have been theoretical exercises (Renton & Poot, 2014; Yoshinaka *et al*., 2018; de Vries *et al*., 2020).

Here, we develop, parameterise, calibrate and validate an E-FSP model of local adaptation along an environmental gradient. As a case study, we use two elevational ecotypes of *Dianthus carthusianorum* that occur along an elevational gradient in the Swiss Alps,growing at low (~1000 m.a.s.l.) and high (~2000 m.a.s.l.) elevation. These environments are characterised by commonly reported differences in (a)biotic conditions along elevational gradients (Halbritter *et al*., 2018), resulting in a tall grassland vegetation at lower elevations, and typical alpine (i.e. shorter) vegetation at higher elevations. Elevational ecotypes of *D. carthusianorum* are adapted to their elevational ranges and display genetically based phenotypic divergence in phenological and morphological traits (Walther, 2020; Pålsson *et al*., *in prep*.). The high elevation populations of *D. carthusianorum* typically exhibit lower biomass, flower earlier and produce smaller flowering stalks compared to their low elevation counterparts. Favoured by a higher energy input environment, the latter achieve larger plant size and taller flowering stalks to potentially compete for light and pollinators with the surrounding vegetation. Despite sound evidence of adaptation along an elevational gradient, the selection pressures underlying the evolution of these elevational ecotypes remains unknown. Commonly reported patterns of adaptation along elevational gradients suggest that the divergence in *D. carthusianorum* is driven by more stressful abiotic conditions at high elevations (e.g. temperature), and by biotic interactions (e.g. competition and pollination) at low elevations (Halbritter *et al*., 2018). First, we aim to validate the E-FSP model by asking whether the E-FSP model can recreate elevational ecotypes of *D. carthusianorum*. Second, we hypothesise that temperature, competition and pollination are key agents of selection and use the E-FSP model to disentangle their relative contribution to elevational adaptation in *D. carthusianorum*.

## Methods

### Model species: D. carthusianorum

*D. carthusianorum* is a primarily outcrossing gynodioecious, perennial herb that is native to Europe and is widespread on rocky slopes and dry grasslands throughout the Alps up to an elevation of 2500 meters (Bloch *et al*., 2006). For model parameterisation, calibration and validation, we used data from two elevational ecotypes of *D. carthusianorum*. These were grown in a reciprocal transplant experiment established in fall 2015 that included two replicate transplant sites at low (~1000 m.a.s.l.) and high (~2000 m.a.s.l.) elevation, respectively. We used data collected in the first growing season on fitness components (survival, flowering probability and seed count), morphological (stalks height, number of stalks, stalk leaves and flowers) and phenological (flowering time) traits (for details see: Walther, 2020; Pålsson *et al*., *in prep*.).

### Model summary

The model used in this study is based on the E-FSP model described in de Vries *et al*. (2020), which was developed in the modelling platform GroIMP (Hemmerling *et al*., 2008) and designed to simulate adaptation to abiotic (nitrogen) and biotic (competition and herbivory) agents. Here, we expand this E-FSP model by including temperature driven plant phenology and plant-pollinator interactions. The model simulates a population of competing plants over multiple generations, with the performance of individual plants within a generation being determined by three plant traits that are subject to selection: germination time (*GM*), time to flowering (*TF*), and stalk height (*SH*). We assumed that these traits are not genetically linked so that there are no pleiotropic effects between them, and therefore the model is theoretically able to select for any combination of trait values. We simulate three environmental factors that determine plant fitness and thereby impose selection pressure; i) the difference in abiotic conditions associated with an elevational gradient (i.e. temperature and subsequently also season length and nitrogen availability), ii) interspecific competition with the surrounding vegetation,and iii) pollinator density. The model structure is summarised below (also see Fig. 1). For a detailed model description, see Methods S1, which includes a list of indices used in the model description (Table S1).

**Fig 1.**
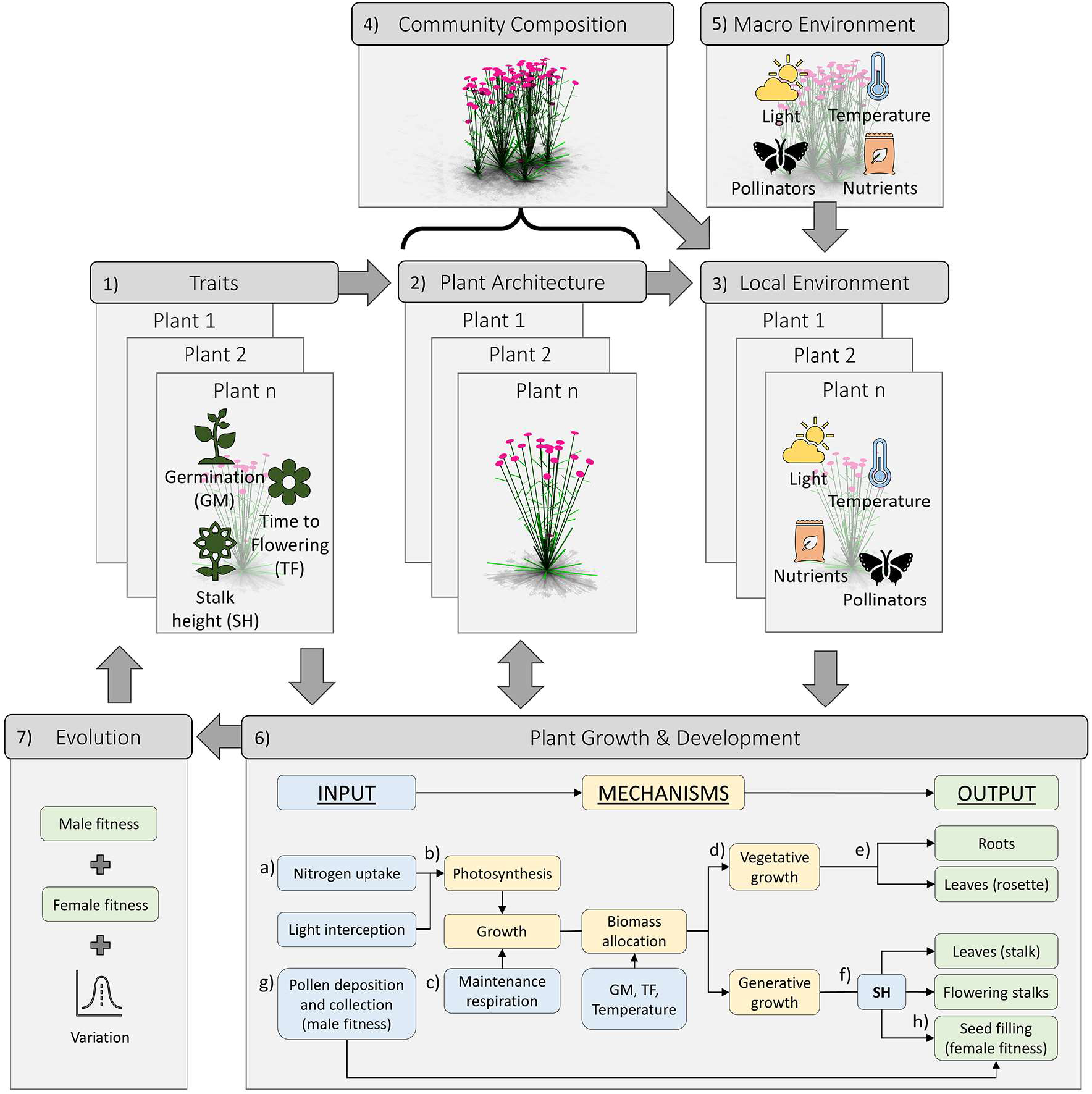
Visual summary of the E-FSP model used in this study. The model represents a population of individual plants, each with their own trait values (**1**; Germination (GM); Time to Flowering (TF); Stalk Height (SH)), plant architecture (**2**) and local environment (**3**; Light, nutrients, pollinators and temperature). The local environment is dependent on the composition of the surrounding plant community (**4**; i.e. intra- and interspecific competition), and the macro environment (**5**). To simulate plant growth and development (**6**), the model takes input on the level of the individual plants (i.e. from **1**,**2** and **3**). First, light interception and nutrient uptake (**a)**are used to calculate leaf level photosynthesis (**b**). The respiration required to maintain the standing biomass (**c**) is subtracted to get the net growth rate. These assimilates are allocated to either vegetative or generative growth (**d**), dependent on the temperature and the GM and TF traits. During vegetative growth, the plant allocates assimilates to roots and rosette leaves (**e**). During generative growth, the plant allocates assimilates to stalk leaves, flowering stalks and seed filling, with the SH trait determining the allocation between these three (**f**). Finally, through recombination, the pollen collected by pollinators (**g**; male fitness) and the filled seeds from pollinated flowers (**h**; female fitness) determine the traits of the plants in next generation, and thus drive evolution through natural selection (**7**).

### Temperature and plant development

The model calculates daily average and minimum temperature as a function of elevation, based on climate data collected by weather stations at the field sites (Fig. S1, Table S2). Temperature is used to drive plant phenology (McMaster & Wilhelm, 1997), photosynthesis (Farquhar *et al*., 1980), soil nitrogen availability (Rodrigo *et al*., 1997; Kirschbaum, 2000; Guntiñas *et al*., 2012), and to calculate frost damage (Ji *et al*., 2015).

Plant development is split into two stages; a vegetative and a generative stage. The *in silico* plants germinate in spring, the timing of which is a function of temperature and their germination trait (GM). During its vegetative stage, the plant invests all accumulated assimilates and nitrogen towards the growth of rosette leaves and roots. The transition to the generative stage is dependent on both the time to flowering trait (TF), and cumulative temperature time (growing degree days, McMaster & Wilhelm, 1997). In the generative stage, the plant continues to intercept light and produce assimilates through photosynthesis, but no longer grows new rosette leaves or roots. Instead, newly acquired assimilates and nitrogen are allocated to flowering stalks, stalk leaves and seed filling.

### Plant architecture and resource capture

The model uses an explicit description of plant architecture to mechanistically simulate competition for the three resources incorporated in the model; light, nitrogen and pollinators. The vegetative shoot is represented by a rosette of rectangular leaves (Fig. S5, Fig. S8) that photosynthesise based on leaf level light interception. The flowering stalks are described as a cylinder with a number of short stalk leaves that also add to assimilate production through photosynthesis and a disk at the top of the stalk that represents the flowerhead and attracts pollinators. The stalk height trait determines the position of the flower, the number of stalk leaves and the stalk diameter required to support the stalk (Fig. S7). The explicit representation of these aboveground plant parts allows for the calculation of light interception in the canopy using a Monte-Carlo ray tracer and, therefore, the outcome of competition for light between individual plants (Hemmerling *et al*., 2008; Evers *et al*., 2010). This methodology has proven to capture the asymmetry in competition for light (de Vries *et al*., 2018; de Vries *et al*., 2019), making it a key model component to simulate the effect of competitive interactions on plant fitness and subsequent selection. The root architecture is described as a conical volume (Fig. S2, Table S3) from which the plant can take up nitrogen, such that a larger root system proportionally increases the potential nitrogen uptake of the plant, thus resulting in symmetric competition for nitrogen.

### Plant growth

We assumed that the C:N ratio of plant tissue is conserved, so that plant growth is either limited by the plant’s ability to intercept light and assimilate carbon through photosynthesis, or its ability to take up nitrogen through the root system. Photosynthesis is calculated at the leaf level using a temperature driven Farquhar, von Caemmerer and Berry photosynthesis model (Farquhar *et al*., 1980; Yin & Struik, 2009; Yin *et al*., 2009). The assimilates produced by the leaves are first used to pay for maintenance respiration, which is based on plant nitrogen content (Ryan, 1991), after which the remaining assimilates are allocated to growth. The potential nitrogen uptake by the root system is modelled as a function of rooting volume and soil nitrogen availability, and can be supplemented by re-allocation of nitrogen from the leaves, which is used to simulate leaf senescence at the end of the growing season (Yin & van Laar, 2005). In the vegetative stage, assimilates and nitrogen are equally allocated towards the growth of rosette leaves and roots (i.e. assuming a root:leaf ratio of 1). In the generative stage, assimilates and nitrogen are allocated to flowering stalks and seed set using a hierarchical allocation model that prioritises filling pollinated seeds (see *Pollination* below) over growing new stalks (Minchin & Thorpe, 1996).

### Pollination

The number of pollinator visits is simulated as a function of flower density following the correlation found by Richman *et al*. (2020). Pollination in grasslands is known to scale with stalk height in relation to the height of the surrounding vegetation (Sletvold *et al*., 2013; Slaviero *et al*., 2016). To simulate this, we visualise the flower heads as upwards facing disks with a diameter of 2 cm, and use the light absorbed by the flowerhead as a proxy for flower attractiveness so that flowers reaching to the top of the vegetation are the most attractive to pollinators. The pollinator visits are then distributed over the flowers in the plot based on their relative attractiveness. The relationship between the number of pollen visitations and potential seed set is based on a previous study of the pollination of *D. carthusianorum* that was conducted in the same study area (Bloch *et al*., 2006, also see Table S4 and Fig. S4).

### Evolutionary algorithm

*D. carthusianorum* is a perennial species that has a lifespan from one to several years. However, implementation of the perennial life-cycle in the model has proven challenging because we currently lack long-term data for calibration and validation. Although life history traits linked to the perennial life-cycle almost certainly contribute to adaptation in *D. carthusianorum*, phenotypic divergence and the resulting differences in performance are already apparent in the first year (Pålsson *et al*., *in prep*.). Therefore, we opted to simulate an annual life-cycle and assume that selection acting on the first year of plant reproduction is sufficient to explain the evolution of the traits under investigation.

Plant fitness is composed of female and male reproductive success: female reproductive success is defined as the total number of filled seeds at the end of the growing season (i.e. fecundity), and male reproductive success is defined as the number of pollinator visits to the flowers over the course of the growing season. At the end of every generation, the model randomly selects 100 plants based on their realised female reproductive success, and 100 plants based on their male reproductive success, and recombines these to generate the 100 offspring genotypes that populate the subsequent generation of plants. In this process, the model allows for a single plant to contribute multiple seeds and/or pollen to the next generation, and we assume that seeds can only germinate in the following year, so no seed bank is built up. The virtual plants were not able to self-pollinate, as selfing in *D. carthusianorum* is prevented by protandry and rarely leads to fruit formation (Bloch *et al*., 2006). The traits of an offspring plant (*T_o_*, dimensionless, ranging from zero to one) are randomly inherited from either of the two parental plants, effectively simulating a haploid system with traits completely determined by their genetic basis. Offspring trait values are assumed to be normally distributed around the parental trait value (mean = *T_p_*, dimensionless, ranging from zero to one, standard deviation = *Tsd*).

### Plant density and interspecific competition

The model starts each generation with 100 individuals of the virtual *D. carthusianorum* plants that were randomly placed in a plot of 1 m^2^, meaning that individual plants can experience different levels of competition dependent on neighbour proximity. While initial plot level plant density is kept constant, the model does allow plant density to vary during the season as a result of mortality caused by frost damage or resource limitation.

To simulate the effect of interspecific competition for light and nutrients, particularly with the tall grasses that *D. carthusianorum* typically competes with in their low elevation habitats, we introduce 100 individuals of a second plant species designed to represent these tall grasses. This increases the initial plant density of the population from 100 plants m^−2^ in the absence of the grass species to 200 plants m^−2^ in its presence. The growth and development of these grasses is not simulated mechanistically, but rather described by a sigmoid function that calculates biomass as a function of temperature and time (see Methods S1). The grasses can therefore be seen as a static environmental factor that imposes competition pressure on the virtual *D. carthusianorum* plants, but is not affected by the *D. carthusianorum* plants.

### Model output

A single simulation consists of 125 generations, by which time the simulated population had settled at an optimum through natural selection (Fig. S9). To account for random fluctuations between generations, model output was recorded at generation 105, 110, 115, 120 and 125. Model output was recorded at the end of the growing season on the level of individual plants and consisted of values for the three plant traits under selection, as well as flowering time, rosette area, the number of stalks, and fitness. We conducted no statistical analyses on the *in silico* data, because the sample size is so high that all treatment combinations show a high statistical significance, even if the differences between those treatment combinations are not biologically relevant. In the text, values are reported as mean ± standard error.

### Model parameterisation, calibration and validation

To parameterise, calibrate and validate the model, we used data collected during the first growing season (2016) of the field experiment described in Pålsson *et al*. (*in prep*.) and conducted a climate chamber experiment to measure germination times. For model parameterisation, we obtained empirical estimates of parameter values that are not readily available in published literature and that we assume to be shared between the elevational ecotypes (for a full list of parameters, see Table S4). For model validation of traits that differentiate the two elevational ecotypes, we obtained empirical estimates growing from a controlled environment (a common garden at the low elevation site for TF, SH, and two climate chambers for GM). Additionally, we obtained empirical estimates of response variables (flowering time, number of stalks, rosette area, and fitness) from the two elevational ecotypes growing in the field. For the germination experiment, we collected seeds from seven individuals from a low and a high elevation population occurring in close proximity (<1km) to the transplant sites. We vernalized the seeds for a week at −18 °C and sowed 20 seeds from each individual in compartmentalised trays with 21 cm^3^ soil per compartment, sowing one seed per compartment. These trays were placed in one of two climate chambers under a constant temperature of either 4 °C or 20 °C to measure the effect of temperature on germination rates, and a 16/8 day/night cycle. We followed a balanced randomised design for this experiment, so that each seed family was equally represented in each treatment. Thrice weekly, the seeds were watered and germination was recorded over a period of 30 days.

### Simulations

The model incorporated three environmental factors; the difference in abiotic conditions associated with a change in **elevation** (i.e. temperature and subsequently also season length and nitrogen availability), interspecific competition with a tall grass species (hereafter named “**competition**”), and pollinator density (hereafter named “**pollination**”, see Table 1). The elevation treatments are 1000 or 2000 m, based on the elevation of the experimental sites. The competition treatments reflect the more intense competition that is generally found in low elevation habitats (Halbritter *et al*., 2018) by simulating both intra- and interspecific competition (i.e. 100 plants m^−2^ of *D. carthusianorum* and 100 plants m^−2^ of a tall grass), and the shorter vegetation that is generally found in high elevation habitats (Halbritter *et al*., 2018) by simulating only intraspecific competition (i.e. only 100 plants m^−2^ of *D. carthusianorum*). The pollination treatments represent pollinator densities along an elevational gradient in the Swiss Alps (Richman *et al*., 2020), with pollinators being more abundant in the low elevation habitat (0.3 pollinator visits flower^−1^ h^−1^) compared to the high elevation habitat (0.03 visits flower^−1^ h^−1^).

To validate model performance, we simulated selection in two scenarios that represent the low and high elevation habitats (**Low habitat**: 1000m elevation, 100 interspecific competitors m^−2^, and 0.3 pollinator visit flower^−1^ h^−1^; **High habitat**: 2000m elevation, 0 interspecific competitors m^−2^, and 0.03 pollinator visit flower^−1^ h^−1^), and compared the trait variation and performance of *in silico* populations after 125 generations of selection to the trait variation and performance of *in vivo* ecotypes of *D. carthusianorum* from the low and high elevation sites. To test whether the *in silico* populations of *D. carthusianorum* could be considered locally adapted, we simulated a virtual transplant experiment: 50-50 mixtures consisting of plants originating from the low and high elevation populations were grown in the low and high elevation habitats, and their seed production after one generation was used as a fitness proxy. Under the basic principles of testing adaptation in reciprocal transplant experiments, genotype x environment (GxE) interactions should result in locally adapted populations outperforming foreign populations growing under the same conditions (Blanquart *et al*., 2013; Hargreaves & Eckert, 2019; Hargreaves *et al*., 2020).

To elucidate how the different abiotic and biotic selection pressures contributed to the local adaptation of *D. carthusianorum*, we first simulated populations under control conditions (i.e. 1000 m, 0 interspecific competitors m^−2^, 0.3 pollinator visit flower^−1^ h^−1^), and then changed each environmental factor individually to assess their effects on trait variation and performance.

## Results

### Simulation of elevational ecotypes: growing in a common garden

To validate whether the model was able to recreate the elevational ecotypes of *D. carthusianorum*, we compared the trait values of *in silico* populations to *in vivo* measurements of the low and high elevation ecotypes conducted on plants growing in a shared environment (i.e. the climate chamber for GM, and the low elevation habitat for SH and TF). The *in vivo* low and high elevation populations of *D. carthusianorum* expressed significant differences in each of the tree selected traits (Fig. 2a; Table S5; *in vivo*). Growing in the climate chamber, plants from the high elevation population germinated later compared to plants from the low elevation population (P<0.001; Fig. S6). Growing in the low elevation site, plants from the high elevation population had a shorter stalk height (P<0.001), and a shorter time to flowering (P<0.001) compared to plants from the low elevation population. The *in silico* populations showed patterns of selection that were equal to the qualitative differences in the *in vivo* populations (Fig. 2a; Table S5).

**Fig. 2.**
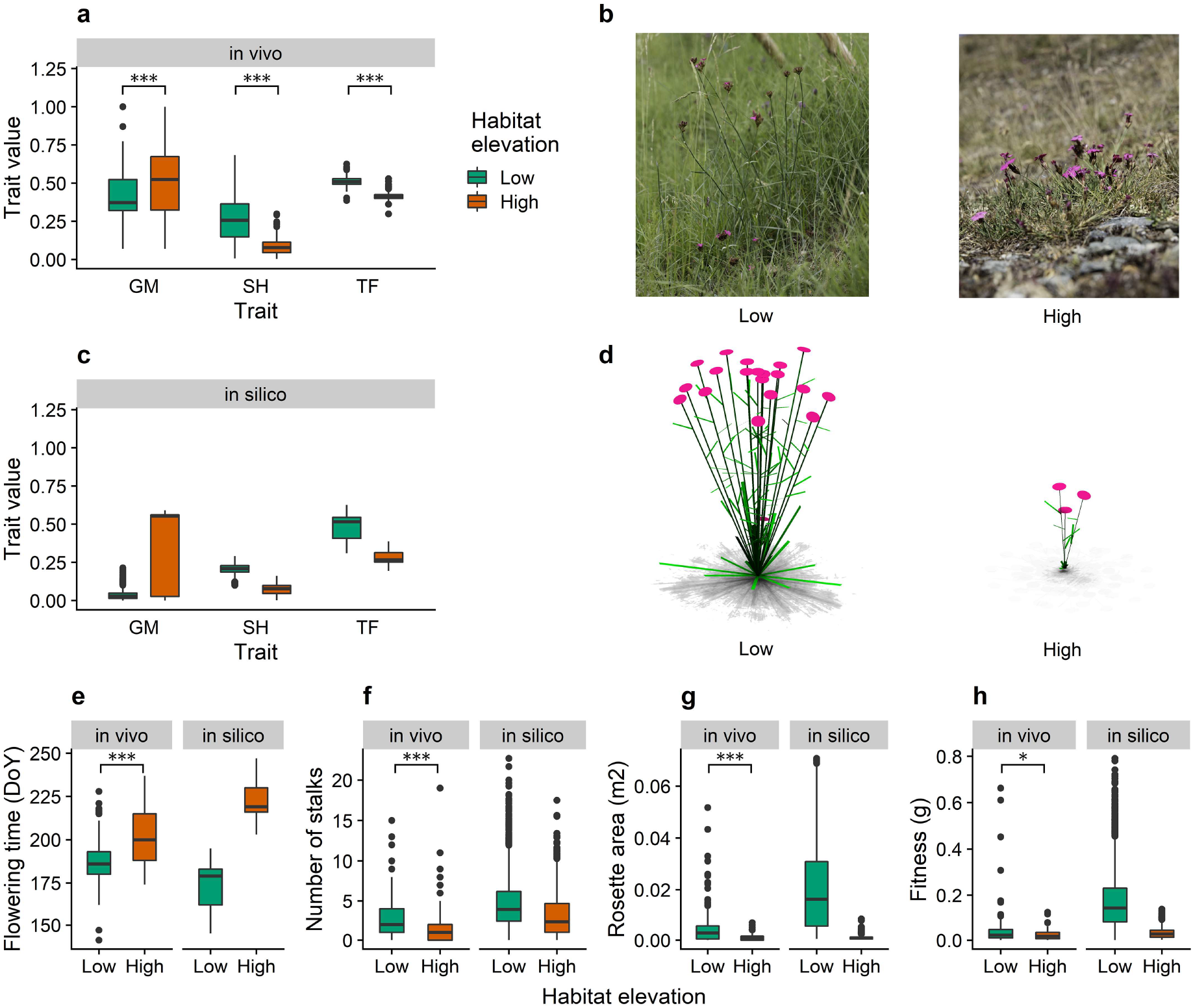
Comparison of trait variation and performance of *in vivo* and *in silico* populations of *D. carthusianorum* in their low and high elevation habitats (Low: Green, High: Red). Trait variation (y-axis: normalized trait value (0-1)) of germination (GM), stalk height (SH) and time to flowering (TF) of *in vivo* (**a,b**) and *in silico* (**c,d**) populations subjected to selection in the low and high elevation habitats. The other panels show the variation in flowering time (**e**, DoY), number of stalks (**f**), rosette area (**g**, m^2^) and fitness (**h**, g) of *in vivo* and *in silico* populations growing in their home environments. Significance is shown only for the measured data (* P<0.05; ** P<0.01; *** P<0.001).

### Simulation of elevational ecotypes: growing in their home environment

To validate the model’s ability to capture GxE interactions, we compared the performance of *in vivo* and *in silico* populations using four performance measures that are determined by both the plant’s trait values and its local environment. From here onwards, we will use the term ‘home environment’ in relation to a plant population to refer to the environment in which selection took place. Growing in their respective home environments, the *in vivo* results show that plants from the high elevation populations flowered significantly later (Fig. 2e; Table S5; P<0.001), produced fewer flowering stalks (Fig. 2f; Table S5; P<0.001), a smaller rosette area(Fig. 2g; Table S5; P<0.001), and lower seed production (Fig. 2h; Table S5; P<0.001) compared to plants from the low elevation populations. Again, the *in silico* results matched the qualitative patterns of the *in vivo* results (Fig. 2b,c,d,e; Table S5).

### Simulation of elevational ecotypes: local adaptation

To test GxE interactions indicative of adaptation in the *in silico* plants, we grew 50:50 mixtures of low and high elevation genotypes under alternative environments. Plants growing in their home environment were able to outcompete the plants originating from the foreign population (Fig. 3). In the low elevation habitat, the plants originating from the low elevation habitat produced more seeds (0.197±0.007 g) than the plants originating from the high elevation habitat (0.026±0.045 g). In the high elevation habitat, the performance of the plants originating from the low elevation habitat saw a major decrease, resulting in them producing fewer seeds (0.008±0.0015 g) than the plants originating from the high elevation habitat (0.039±00026 g). These results fulfil both the local vs foreign and home vs away criteria forming the hallmarks of local adaptation (Kawecki & Ebert, 2004; Savolainen *et al*., 2013).

**Fig. 3.**
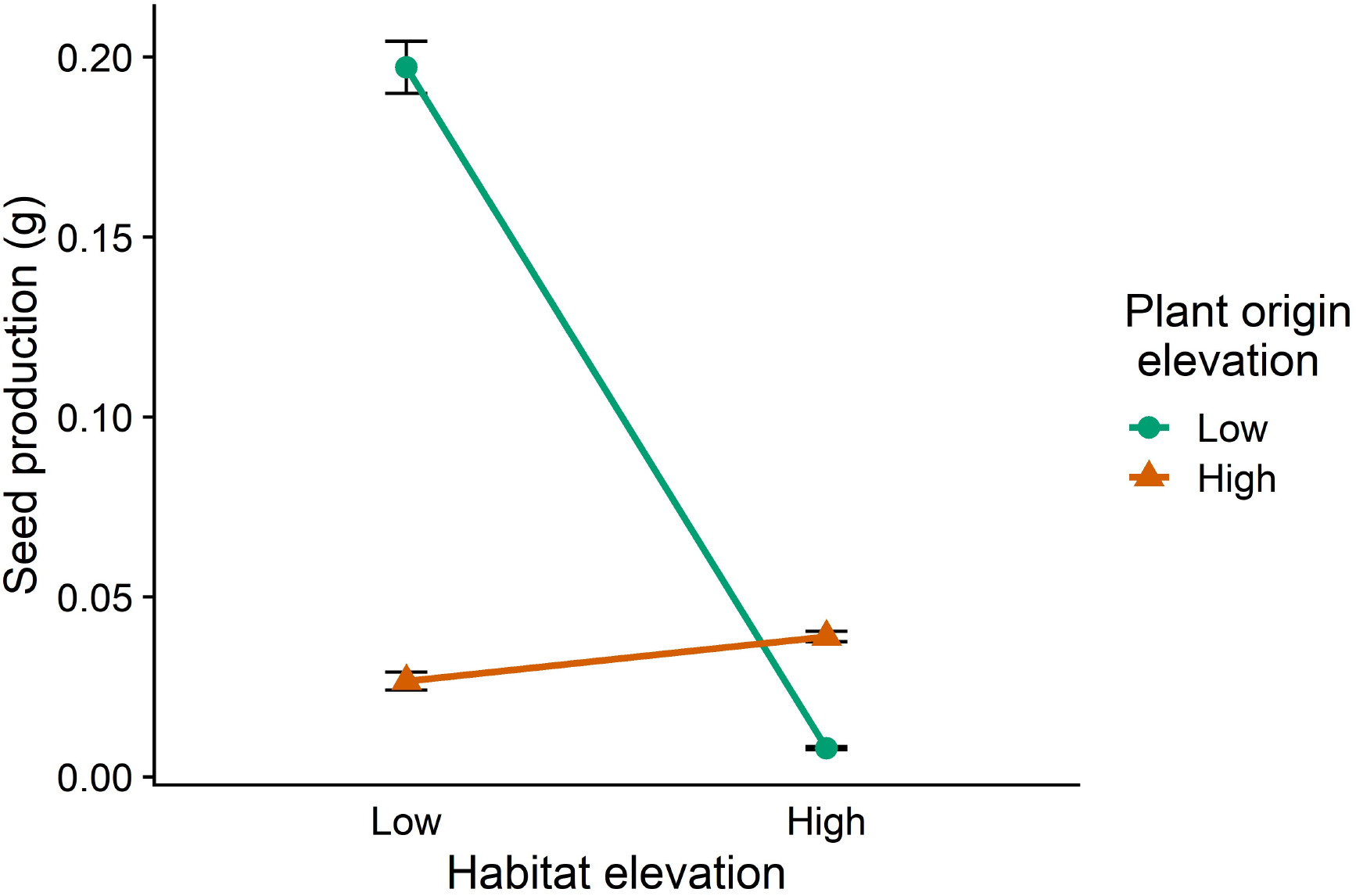
Seed production of a virtual transplant experiment in the low and high elevation habitats. We simulated plant populations consisting of a 50:50 mixture of plants originating from the *in silico* low (green) and high (red) elevation habitats competing in either the low and high elevation habitats. Error bars show the standard error of the mean.

### Disentangling the role of individual selection pressures: Elevation

Increasing elevation from 1000m to 2000m decreased daily average temperature by ~5°C, which decreased the season length by 108 days (assuming a base temperature of zero) and decreased nitrogen mineralisation over the year from 329 g N m^−2^ to 182 g N m^−2^. Additionally, this increase in elevation increased the variation in minimum temperature (eq. S30), increasing the frequency and strength of freezing events that potentially lead to frost damage. These changes in the environment led to selection for plants that germinated later compared to the control treatment (Fig. 4a, GM), which allowed the plants to escape the increased risk of frost damage early in the season. These environmental changes also selected for shorter flowering stalks (Fig. 4a, SH), and earlier flowering time compared to the control treatment (Fig. 4a, TF). These changes in plant traits led to plants that, following adaptation to high elevation and grown in a control environment, flowered earlier compared to plants from the control treatment (Fig. 4b), and also produced fewer stalks (Fig. 4c), a smaller rosette (Fig. 4d) and lower fitness (Fig. 4e). However, when growing in their home environment, these plants still flowered later than the control plants in the control treatment because of the late start of the season at high elevation (Fig. 4f). The decrease in temperature associated with the increase in elevation led to a decrease in productivity through a decrease in photosynthetic rates, a shorter growing season and lower nitrogen availability. This lower productivity in combination with the trait changes led to the plants having smaller rosettes (Fig. 4g), fewer flowering stalks (Fig. 4h), and lower seed production (Fig. 4i).

**Fig. 4.**
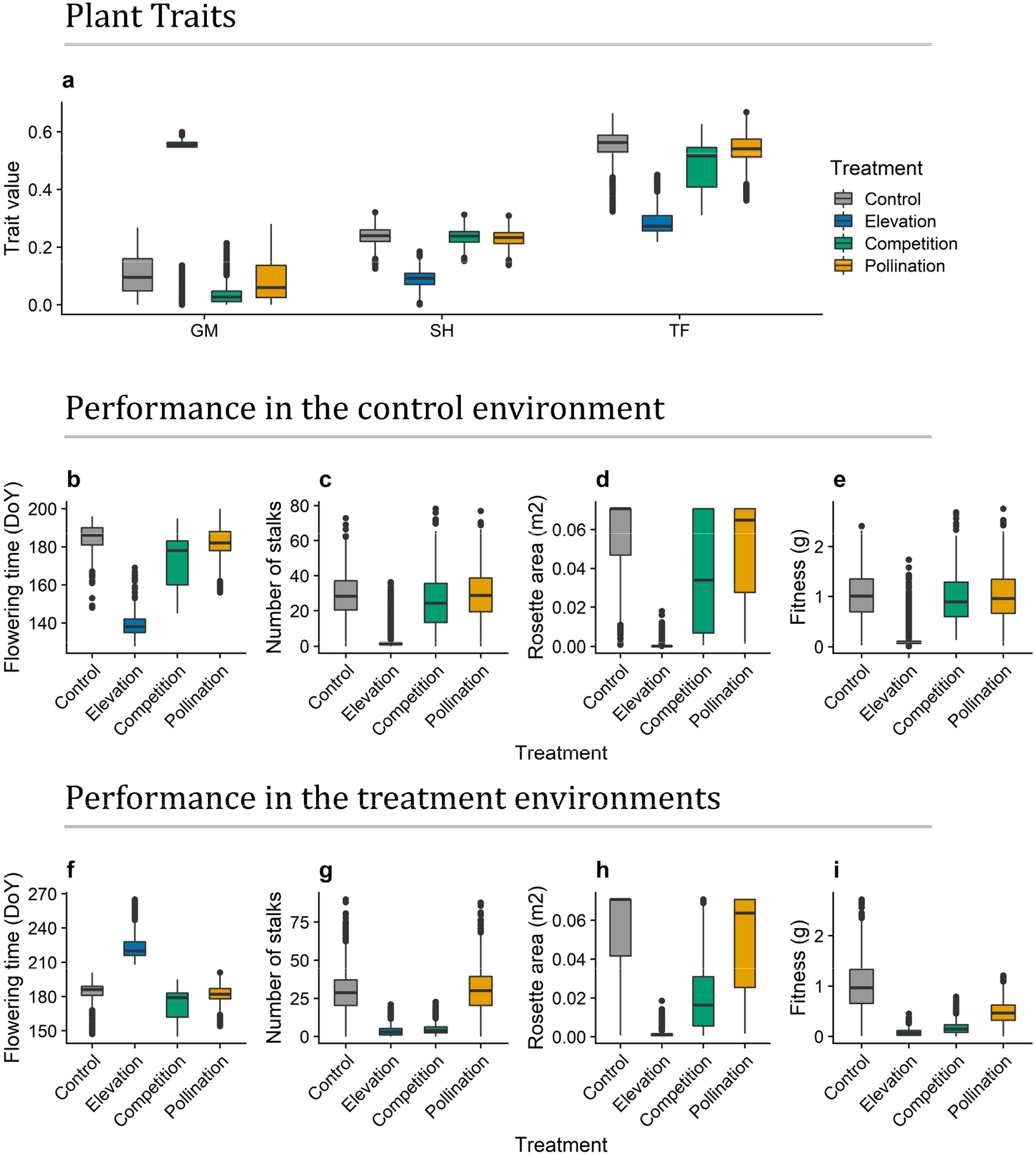
Trait selection on *in silico* populations of *D. carthusianorum*. Panel **a** shows the trait values of three plant traits (**a**; normalized trait value (0-1); germination (GM), stalk height (SH) and time to flowering (TF)) resulting from selection imposed by treatments were we varied individual environmental factors (**Control**, grey; changes in abiotic conditions associated with an increased **Elevation**, blue; increased **Competition**, green; or decreased **Pollination**, yellow). Panels **b-e** show the performance of the adapted plant populations in the control environment to show the direct effect of the trait changes on plant performance. Panels **f-i** show the performance of the adapted plant populations in their respective home environments (i.e. the environment in which selection took place), which shows the combined effect of changes in environment and plant traits on plant performance.

### Disentangling the role of individual selection pressures: Interspecific competition

In the competition treatment, the *in silico D. carthusianorum* plants competed for light and nitrogen with an equal density of a tall grass species. This interspecific competition selected for earlier germination (Fig. 4a, GM), and a small decrease in the time to flowering (Fig. 4a, TF), but not for a change in stalk height compared to the control treatment (Fig. 4a, SH). Growing in the control environment, these trait changes resulted in a slightly earlier flowering time (Fig. 4b) and small decreases in the number of stalks (Fig. 4c), the rosette area (Fig. 4d), and fitness (Fig. 4e). In their high competition home environment, the plants flowered slightly earlier (Fig. 4f), and increased competition led to a major reduction in productivity, resulting in major decreases in the number of stalks (Fig. 4g), rosette area (Fig. 4h), and fitness (Fig. 4i) compared to the plants growing in the control treatment.

### Disentangling the role of individual selection pressures: Pollination

In the pollination treatment, the decrease in pollinator abundance led to a shift from seed production being mostly carbon limited, to seed production being more pollen limited and an increase in unfilled seeds (Fig. S10). This shift in limitation leads to a decrease in fitness (Fig. 4i) without a decrease in productivity, as the number of stalks (Fig. 4g) and rosette area (Fig. 4h) did not change relative to the control treatment. The decrease in pollinator density selected for slightly earlier germination (Fig. 4a, GM), but there were no changes in selection for stalk height and time to flowering (Fig. 4a, SH, TF). This resulted in plants that flowered slightly earlier compared to plants from the control treatment (Fig. 4b), but achieved an equal number of stalks (Fig. 4c), rosette area (Fig. 4d) and fitness (Fig. 4e) in the control environment.

## Discussion

### Simulation of elevational ecotypes

Adaptation to local conditions is a key mechanism in the evolution and diversification of plant species (Hargreaves & Eckert, 2019; Hargreaves *et al*., 2020). Our E-FSP model was able to reproduce the patterns of local adaptation along an elevational gradient found in *D. carthusianorum*. The model reproduced the qualitative differences between two elevational ecotypes in two phenological (germination and time to flowering) and one morphological trait (stalk height), as well as qualitative differences in four variables related to plant performance that emerge from GxE interactions (flowering time, number of stalks, rosette area and seed production). Moreover, the model satisfied the home vs away and local vs foreign criteria that indicate populations are locally adapted to their home environments, in line with empirical evidence (Pålsson *et al*., *in prep*.). It is remarkable that the model was able to recreate these patterns of local adaptation in a complex natural system where selection is driven by multiple abiotic and biotic agents.

So far, FSP models have mostly focussed on agricultural (Lopez *et al*., 2010; Zhu *et al*., 2015; Evers & Bastiaans, 2016; Coussement *et al*., 2020), horticultural (Sarlikioti *et al*., 2011; Chen *et al*., 2014; Dieleman *et al*., 2019; Zhang *et al*., 2020), and model systems (Bongers *et al*., 2018). FSP models that simulate natural systems with an increased ecological complexity are seeing recent development, yet these models are still being validated on data collected under controlled experimental conditions (de Vries *et al*., 2018; Faverjon *et al*., 2019) and still lack the ecological variability and complexity that shape plant communities (Bongers, 2020; de Vries, 2021). Here, we validated our model on data from a transplant experiment where plants grew under natural conditions, which, to our knowledge, is the first time an E-FSP model has been validated to empirical data collected under natural conditions. The model’s ability to recreate the patterns of selection exerted by such a complex environment highlights the potential of this approach for learning more about selection and to study the complex eco-evolutionary dynamics that shape natural plant communities.

### Disentangling the role of individual selection pressures: the abiotic environment

Our results suggest that in the case of *D. carthusianorum*, the abiotic environment is the most important driver of elevational adaptation, imposing strong selection pressure on both the phenological (germination and flowering times) and morphological (stalk height) traits. These selection pressures resulted in high elevation plants that, growing in a shared environment following adaptation, flowered earlier, were shorter and accumulated less biomass than plants from the low elevation population. These results match commonly reported trends in studies of plant adaptation along elevational gradients (Halbritter *et al*., 2018). Evolution towards smaller size is generally assumed to be advantageous in alpine environments due to warmer microclimates close to the ground, increased protection from wind, or a result of selection for increased stress resistance (Körner, 2003). Interestingly, our model did not implement microclimate, wind or stress resistance, yet reproduced this pattern of adaptation through divergence in phenological traits. High elevation genotypes germinate late to avoid frost damage and flower fast to complete the reproductive cycle within the summer season, resulting in a plant phenotype that has a shorter vegetative stage. This shorter period of vegetative growth leads to lower potential for biomass accumulation, a decrease in the number of stalks and a decrease in reproductive fitness. Thus, the shift in phenology may be responsible for lower stalk height, because of the decrease in potential biomass accumulation. Overall, our results suggest that divergence in phenological traits potentially compound direct selection pressure for shorter and smaller phenotypes at high elevations.

### Disentangling the role of individual selection pressures: the biotic environment

In our model, both interspecific competition and decreased pollination strongly decreased plant fitness, but contributed comparatively little to local adaptation. This is in concordance with the findings of a recent meta-analysis that showed biotic interactions generally do not make patterns of local adaption stronger or more common (Hargreaves *et al*., 2020), despite having a strong and well documented effect on plant performance (e.g.Weiner, 1990).

In our model, interspecific competition selected for earlier germination and earlier flowering, increasing the resource capture early in the season when interspecific competition was lower, but also increasing the risk of frost damage early in the season. This highlights how trade-offs between different components of plant performance (e.g. biomass accumulation and survival) can drive selection in opposite directions, potentially resulting in stabilising selection. Surprisingly, interspecific competition did not select for an increase in stalk height compared to the control treatment (i.e. Control vs Competition, see Fig. 4a). This contradicts expectations, as increased height is a well-known response of plants that grow in a competitive environment (Ballaré *et al*., 1990; Falster & Westoby, 2003). Traits such as leaf angle, leaf shape and especially stem elongation are known to be key determinants of the outcome of competition for light as they determine leaf light interception by mediating the position of leaves relative to the surrounding vegetation (Franklin, 2008; Ballaré & Pierik, 2017). However, plants growing in competition can be equally tall as plants growing in the absence of competition, yet with a much higher height to biomass ratio caused by decreased biomass accumulation under competition, making the same investment in height growth relatively more costly (de Vries *et al*., 2018). When considering the investment in height relative to plant biomass, both the *in vivo* and *in silico* populations of *D. carthusianorum* growing with interspecific competition show the increased investment in height growth that is expected in a competitive environment.

In our model, reduced pollinator abundance decreases plant fitness, but does not affect selection. The *in silico* plants can increase their competitiveness for pollinators in one of three ways; produce more flowers, produce taller stalks, or have a longer generative stage of development and thus have a longer period in which to attract pollinators. While each of these traits will increase male fitness and potential female fitness, they also severely restrict the plant’s ability to accumulate biomass, compete for light and nutrients, and to fill seeds. The model did not include flower traits, which are known to be strong drivers of pollinators visitation (Fornoff *et al*., 2017; Walther, 2020), and are known to differ along elevational gradients (Fabbro & Körner, 2004). Flower traits may be the main mechanism for plants to increase pollinator attraction as they potentially come at lower (opportunity) costs than the mechanisms included in our model.

### Future model development

Here, we have shown the potential for E-FSP modelling to simulate the emergent behaviour of a complex natural system that includes abiotic and biotic agents, and integrates physiological, ecological and evolutionary mechanisms. E-FSP modelling is a promising and versatile tool that is capable of simulating more complex and dynamic systems than is common in FSP modelling, and integrates more physiological and spatial detail than commonly used eco- evolutionary modelling approaches. We would like to highlight two avenues of future model development for E-FSP models: simulation of multi-species communities with different life-history strategies, and simulation of complex genetic and demographic processes that shape local adaptation.

FSP modelling has proven to be capable of simulating the growth and development of a wide range of plant species (Dunbabin *et al*., 2013; Pagès *et al*., 2014; Louarn & Song, 2020), and FSP modelling is being used to simulate multi-species systems in an agricultural setting (Evers *et al*., 2019). Conversely, FSP models that focus on natural systems are often used to simulate single plant species rather than diverse mixed-species communities, which have received only recent attention (Faverjon *et al*., 2019; Bongers, 2020; de Vries, 2021). Here, we have focussed on a single plant species, but have shown the model’s ability to simulate the diversifying forces of selection. A key point of focus for the future development of E-FSP models is the simulation of different life-history traits and species co-existence. By simulating a community consisting of multiple species, the model can theoretically select for different life-history traits that fill different niches. The main challenge for the implementation of different life-history traits lies in the complexity of carbohydrate and nitrogen cycles in perennial plants, which leads to difficulties in linking theory to observations and formulating a comprehensive mechanistic model (Monson *et al*., 2006).

Future development of E-FSP modelling can see the incorporation of more detail in genetic and demographic processes that drive population and community dynamics (Lowe *et al*., 2017). In particular, gene flow between populations is known to play a complex eco-evolutionary role as it can either promote or constrain adaptation, dependent on the migration-selection balance (Garant *et al*., 2007). Gene flow is traditionally seen as a force that homogenises populations by working against the diversifying forces of selection, which drive local adaptation (Haldane, 1930; García-Ramos & Kirkpatrick, 1997). However, recent studies show that local adaptations can be maintained despite high gene flow provided that selection coefficients can sustain ecotypic divergence (Gonzalo-Turpin & Hazard, 2009; Fitzpatrick *et al*., 2015; Tigano & Friesen, 2016; Luqman *et al*., 2021). On the other hand, low amounts of gene flow between locally adapted populations can be beneficial as they allow adaptive alleles to spread across populations and lead to genetic rescue in the face of rapid environmental change (Slatkin, 1987; Rieseberg & Burke, 2001; Tallmon *et al*., 2004). E-FSP models can contribute to our understanding of the role gene flow plays in mediating the responses of plant communities to environmental change, particularly because the strength of selection, and thus the migration-selection balance, emerges naturally from interactions between mechanisms implemented in the FSP model.

The model presented here represents a major advance in the development of mechanistic models that incorporate physiological, ecological and evolutionary mechanisms to simulate the complexity of plant phenotypic variation. We have shown the promise of this methodology to explore the ecological complexity that drives local adaptation in natural plant communities, thereby complementing experimental and statistical modelling approaches. The approach offers a tool to better understand what mechanisms and selective agents drive local adaptation, and how local adaptation mediates the response of plant communities to rapid environmental change.

## References

Ballar CL, Pierik R. 2017. The shade-avoidance syndrome: multiple signals and ecological consequences. Plant, Cell & Environment 40(11): 2530–2543.

Ballar CL, Scopel AL, Sanchez RA. 1990. Far-red radiation reflected from adjacent leaves: an early signal of competition in plant canopies. Science 247(4940): 329–332.

Blanquart F, Kaltz O, Nuismer SL, Gandon S. 2013. A practical guide to measuring local adaptation. Ecology Letters 16(9): 1195–1205.

Bloch D, Werdenberg N, Erhardt A. 2006. Pollination crisis in the butterfly-pollinated wild carnation Dianthus carthusianorum? New Phytologist 169(4): 699–706.

Bongers FJ. 2020. Functional-structural plant models to boost understanding of complementarity in light capture and use in mixed-species forests. Basic and Applied Ecology 48: 92–101.

Bongers FJ, Douma JC, Iwasa Y, Pierik R, Evers JB, Anten NP. 2019. Variation in plastic responses to light results from selection in different competitive environments—A game theoretical approach using virtual plants. PLoS computational biology 15(8).

Bongers FJ, Pierik R, Anten NPR, Evers JB. 2018. Subtle variation in shade avoidance responses may have profound consequences for plant competitiveness. Annals of Botany 121(5): 863–873.

Briscoe Runquist RD, Gorton AJ, Yoder JB, Deacon NJ, Grossman JJ, Kothari S, Lyons MP, Sheth SN, Tiffin P, Moeller DA. 2020. Context dependence of local adaptation to abiotic and biotic environments: a quantitative and qualitative synthesis. The American Naturalist 195(3): 412–431.

Chen T-W, Henke M, De Visser PH, Buck-Sorlin G, Wiechers D, Kahlen K, Stützel H. 2014. What is the most prominent factor limiting photosynthesis in different layers of a greenhouse cucumber canopy? Annals of Botany 114(4): 677–688.

Connolly SR, Keith SA, Colwell RK, Rahbek C. 2017. Process, mechanism, and modeling in macroecology. Trends in Ecology & Evolution 32(11): 835–844.

Coussement JR, De Swaef T, Lootens P, Steppe K. 2020. Turgor-driven plant growth applied in a soybean functional–structural plant model. Annals of Botany 126(4): 729–744.

de Vries J. 2021. Using evolutionary functional-structural plant models to understand climate change impacts on plant communities. in silico Plants 3(2).

de Vries J, Evers JB, Dicke M, Poelman EH. 2019. Ecological interactions shape the adaptive value of plant defence: herbivore attack versus competition for light. Functional Ecology 33(1): 129–138.

de Vries J, Evers JB, Poelman EH, Anten NP. 2020. Simulation of optimal defence against herbivores under resource limitation and competition using an evolutionary functional-structural plant model. in silico Plants 2(1).

de Vries J, Poelman EH, Anten NP, Evers JB. 2018. Elucidating the interaction between light competition and herbivore feeding patterns using functional–structural plant modelling. Annals of Botany 121(5): 1019–1031.

Dieleman JA, De Visser PH, Meinen E, Grit JG, Dueck TA. 2019. Integrating morphological and physiological responses of tomato plants to light quality to the crop level by 3D modeling. Frontiers in plant science 10: 839.

Douma JC, de Vries J, Poelman EH, Dicke M, Anten NP, Evers JB. 2019. Ecological significance of light quality in optimizing plant defence. Plant, Cell & Environment 42(3): 1065–1077.

Dunbabin VM, Postma JA, Schnepf A, Pagès L, Javaux M, Wu L, Leitner D, Chen YL, Rengel Z, Diggle AJ. 2013. Modelling root–soil interactions using three–dimensional models of root growth, architecture and function. Plant and Soil 372(1-2): 93–124.

Evers J, Vos J, Yin X, Romero P, Van Der Putten P, Struik P. 2010. Simulation of wheat growth and development based on organ-level photosynthesis and assimilate allocation. Journal of Experimental Botany 61(8): 2203–2216.

Evers JB, Bastiaans L. 2016. Quantifying the effect of crop spatial arrangement on weed suppression using functional-structural plant modelling. Journal of plant research 129(3): 339–351.

Evers JB, Letort V, Renton M, Kang M. 2018. Computational botany: advancing plant science through functional–structural plant modelling. Annals of Botany 121(5): 767–772.

Evers JB, Van Der Werf W, Stomph TJ, Bastiaans L, Anten NP. 2019. Understanding and optimizing species mixtures using functional–structural plant modelling. Journal of Experimental Botany 70(9): 2381–2388.

Fabbro T, Körner C. 2004. Altitudinal differences in flower traits and reproductive allocation. Flora-Morphology, Distribution, Functional Ecology of Plants 199(1): 70–81.

Falster DS, Westoby M. 2003. Plant height and evolutionary games. Trends in Ecology & Evolution 18(7): 337–343.

Farquhar GD, von Caemmerer Sv, Berry J. 1980. A biochemical model of photosynthetic CO 2 assimilation in leaves of C 3 species. Planta 149(1): 78–90.

Faverjon L, Escobar-Gutiérrez A, Litrico I, Julier B, Louarn G. 2019. A generic individual-based model can predict yield, nitrogen content, and species abundance in experimental grassland communities. Journal of Experimental Botany.

Fitzpatrick S, Gerberich J, Kronenberger J, Angeloni L, Funk W. 2015. Locally adapted traits maintained in the face of high gene flow. Ecology Letters 18(1): 37–47.

Fornoff F, Klein AM, Hartig F, Benadi G, Venjakob C, Schaefer HM, Ebeling A. 2017. Functional flower traits and their diversity drive pollinator visitation. Oikos 126(7): 1020–1030.

Franklin KA. 2008. Shade avoidance. New Phytologist 179(4): 930–944.

Garant D, Forde SE, Hendry AP. 2007. The multifarious effects of dispersal and gene flow on contemporary adaptation. Functional Ecology 21(3): 434–443.

García-Ramos G, Kirkpatrick M. 1997. Genetic models of adaptation and gene flow in peripheral populations. Evolution 51(1): 21–28.

Gonzalo-Turpin H, Hazard L. 2009. Local adaptation occurs along altitudinal gradient despite the existence of gene flow in the alpine plant species Festuca eskia. Journal of Ecology 97(4): 742–751.

Guntiñas ME, Leirós M, Trasar-Cepeda C, Gil-Sotres F. 2012. Effects of moisture and temperature on net soil nitrogen mineralization: A laboratory study. European Journal of Soil Biology 48: 73–80.

Halbritter AH, Fior S, Keller I, Billeter R, Edwards PJ, Holderegger R, Karrenberg S, Pluess AR, Widmer A, Alexander JM. 2018. Trait differentiation and adaptation of plants along elevation gradients. Journal of Evolutionary Biology 31(6): 784–800.

Haldane JBS 1930. A mathematical theory of natural and artificial selection.(Part VI, Isolation.). Mathematical Proceedings of the Cambridge Philosophical Society: Cambridge University Press. 220–230.

Hammer G, Cooper M, Tardieu F, Welch S, Walsh B, van Eeuwijk F, Chapman S, Podlich D. 2006. Models for navigating biological complexity in breeding improved crop plants. Trends in Plant Science 11(12): 587–593.

Hargreaves AL, Eckert CG. 2019. Local adaptation primes cold-edge populations for range expansion but not warming-induced range shifts. Ecology Letters 22(1): 78–88.

Hargreaves AL, Germain RM, Bontrager M, Persi J, Angert AL. 2020. Local adaptation to biotic interactions: A meta-analysis across latitudes. The American Naturalist 195(3): 395–411.

Hemmerling R, Kniemeyer O, Lanwert D, Kurth W, Buck-Sorlin G. 2008. The rule-based language XL and the modelling environment GroIMP illustrated with simulated tree competition. Functional Plant Biology 35(9-10): 739–750.

Ji H, Wang Y, Cloix C, Li K, Jenkins GI, Wang S, Shang Z, Shi Y, Yang S, Li X. 2015. The Arabidopsis RCC1 family protein TCF1 regulates freezing tolerance and cold acclimation through modulating lignin biosynthesis. PLoS Genet 11(9): e1005471.

Kawecki TJ, Ebert D. 2004. Conceptual issues in local adaptation. Ecology Letters 7(12): 1225–1241.

Kirschbaum MU. 2000. Will changes in soil organic carbon act as a positive or negative feedback on global warming? Biogeochemistry 48(1): 21–51.

Körner C 2003. Alpine plant life: functional plant ecology of high mountain ecosystems: Springer.

Leimu R, Fischer M. 2008. A meta-analysis of local adaptation in plants. PLoS One 3(12): e4010.

Lopez G, Favreau RR, Smith C, DeJong TM. 2010. L-PEACH: a computer-based model to understand how peach trees grow. HortTechnology 20(6): 983–990.

Louarn G, Song Y. 2020. Two decades of functional–structural plant modelling: now addressing fundamental questions in systems biology and predictive ecology. Annals of Botany 126(4): 501–509.

Lowe WH, Kovach RP, Allendorf FW. 2017. Population genetics and demography unite ecology and evolution. Trends in Ecology & Evolution 32(2): 141–152.

Luqman H, Widmer A, Fior S, Wegmann D. 2021. Identifying loci under selection via explicit demographic models. Molecular Ecology Resources.

McMaster GS, Wilhelm W. 1997. Growing degree-days: one equation, two interpretations. Agricultural and Forest Meteorology 87(4): 291–300.

McNickle GG, Dybzinski R. 2013. Game theory and plant ecology. Ecology Letters 16(4): 545–555.

Minchin P, Thorpe M. 1996. What determines carbon partitioning between competing sinks? Journal of Experimental Botany 47(Special_Issue): 1293–1296.

Monson RK, Rosenstiel TN, Forbis TA, Lipson DA, Jaeger III CH. 2006. Nitrogen and carbon storage in alpine plants. Integrative and Comparative Biology 46(1): 35–48.

Pagès L, Bécel C, Boukcim H, Moreau D, Nguyen C, Voisin A-S. 2014. Calibration and evaluation of ArchiSimple, a simple model of root system architecture. Ecological Modelling(290): 76–84.

Pålsson A, Widmer A, Fior S. in prep. Altitudinal adaptation is mediated by life history traits and adaptive plasticity in an alpine carnation

Paquette A, Hargreaves AL. 2021. Biotic interactions are more often important at species’ warm versus cool range edges. Ecology Letters n/a(n/a).

Renton M, Poot P. 2014. Simulation of the evolution of root water foraging strategies in dry and shallow soils. Annals of Botany 114(4): 763–778.

Richman SK, Levine JM, Stefan L, Johnson CA. 2020. Asynchronous range shifts drive alpine plant–pollinator interactions and reduce plant fitness. Global Change Biology 26(5): 3052–3064.

Rieseberg LH, Burke J. 2001. A genic view of species integration. Journal of Evolutionary Biology 14(6): 883–886.

Rodrigo A, Recous S, Neel C, Mary B. 1997. Modelling temperature and moisture effects on C–N transformations in soils: comparison of nine models. Ecological Modelling 102(2-3): 325–339.

Ryan MG. 1991. Effects of climate change on plant respiration. Ecological applications 1(2): 157–167.

Sarlikioti V, de Visser PH, Buck-Sorlin G, Marcelis L. 2011. How plant architecture affects light absorption and photosynthesis in tomato: towards an ideotype for plant architecture using a functional–structural plant model. Annals of Botany 108(6): 1065–1073.

Savolainen O, Lascoux M, Merilä J. 2013. Ecological genomics of local adaptation. Nature Reviews Genetics 14(11): 807–820.

Slatkin M. 1987. Gene flow and the geographic structure of natural populations. Science 236(4803): 787–792.

Slaviero A, Del Vecchio S, Pierce S, Fantinato E, Buffa G. 2016. Plant community attributes affect dry grassland orchid establishment. Plant Ecology 217(12): 1533–1543.

Sletvold N, Grindeland JM, Ågren J. 2013. Vegetation context influences the strength and targets of pollinator-mediated selection in a deceptive orchid. Ecology 94(6): 1236–1242.

Tallmon DA, Luikart G, Waples RS. 2004. The alluring simplicity and complex reality of genetic rescue. Trends in Ecology & Evolution 19(9): 489–496.

Tigano A, Friesen VL. 2016. Genomics of local adaptation with gene flow. Molecular Ecology 25(10): 2144–2164.

Wadgymar SM, Lowry DB, Gould BA, Byron CN, Mactavish RM, Anderson JT. 2017. Identifying targets and agents of selection: innovative methods to evaluate the processes that contribute to local adaptation. Methods in Ecology and Evolution 8(6): 738–749.

Walther U. 2020. The evolution of floral traits in a heterogeneous environment. ETH Zurich.

Weiner J. 1990. Asymmetric competition in plant populations. Trends in Ecology & Evolution 5(11): 360–364.

Yin X, Struik P. 2009. C3 and C4 photosynthesis models: an overview from the perspective of crop modelling. NJAS-Wageningen Journal of Life Sciences 57(1): 27–38.

Yin X, Struik PC, Romero P, Harbinson J, Evers JB, Van Der Putten PE, Vos J. 2009. Using combined measurements of gas exchange and chlorophyll fluorescence to estimate parameters of a biochemical C3 photosynthesis model: a critical appraisal and a new integrated approach applied to leaves in a wheat (Triticum aestivum) canopy. Plant, Cell & Environment 32(5): 448–464.

Yin X, van Laar HH. 2005. Crop systems dynamics: an ecophysiological simulation model for genotype-by-environment interactions. Wageningen, The Netherlands: Wageningen Academic Pub.

Yoshinaka K, Nagashima H, Yanagita Y, Hikosaka K. 2018. The role of biomass allocation between lamina and petioles in a game of light competition in a dense stand of an annual plant. Annals of Botany 121(5): 1055–1064.

Zhang N, Van Westreenen A, Evers JB, Anten NP, Marcelis LF. 2020. Quantifying the contribution of bent shoots to plant photosynthesis and biomass production of flower shoots in rose (Rosa hybrida) using a functional–structural plant model. Annals of Botany 126(4): 587–599.

Zhu J, van der Werf W, Anten NPR, Vos J, Evers JBC. 2015. The contribution of phenotypic plasticity to complementary light capture in plant mixtures. New Phytologist 207(4): 1213–1222.

