## Supporting information for "Unravelling drivers of local adaptation through Evolutionary Functional-Structural Plant modelling"

### Methods S1: Detailed model description

The FSP model used in this study was implemented in the platform GroImp v1.5 (Hemmerling et al. 2008), and based on the evolutionary FSP model described in de Vries et al. (2020).

**Table S1.** List of indices used in the model description.

| Index | Name | Index | Name |
|---|---|---|---|
| p | Plant | A | Assimilates gained through photosynthesis |
| l | Leaf | G | Assimilates available for growth |
| s | Stalk | L | Assimilates allocated to leaves |
| f | Flower | R | Assimilates allocated to roots |
| | | ST | Assimilates allocated to stalks |
| | | SE | Assimilates allocated to seeds |

*Temperature*

The average temperature (*avgT*, ºC) on a given day of the year (*DoY*, d) is calculated as a function of elevation (*Elev*, m) and five parameters that determine the shape of the temperature curve(*avgYT*, average yearly temperature at 0 m elevation, degrees C; *A0*, base amplitude; *dA*, change in amplitude with a 1000 m increase in elevation; *Ph0*, base phase; *dPh*, change in phase with a 1000m increase in elevation; *dYT*, change in yearly average temperature with a 1000 m increase in elevation).

$$avgT = avgYT + \left(\left(A0 + \frac{dA * Elev}{1000}\right) - avgYT\right) * \sin\left(2\pi * \frac{DoY}{365} + \frac{2\pi}{365} * \left(Ph0 + \frac{dPh * Elev}{1000}\right)\right) + \frac{dYT * Elev}{1000} \qquad \text{(S1)}$$

The variation around the average temperature (*sdT*, degrees C) is also simulated as a function of elevation using a parameter for the standard deviation of temperature at sea level (*sdT0*, degrees C) and a parameter for the change in the standard deviation per 1000m of elevation (*dsdT*, degrees C 1000 $m^{-1}$).

$$sdT = dsT0 + dsdT * \frac{Elev}{1000} \qquad \text{(S2)}$$

We parameterised two temperature curves on temperature data collecting at the low and high elevation field sites. The first curve simulates the average day-time temperature and the second curve simulates the minimum night-time temperature (Fig. S1; for parameter values see Table S2), which are used to model plant growth and frost damage, respectively.

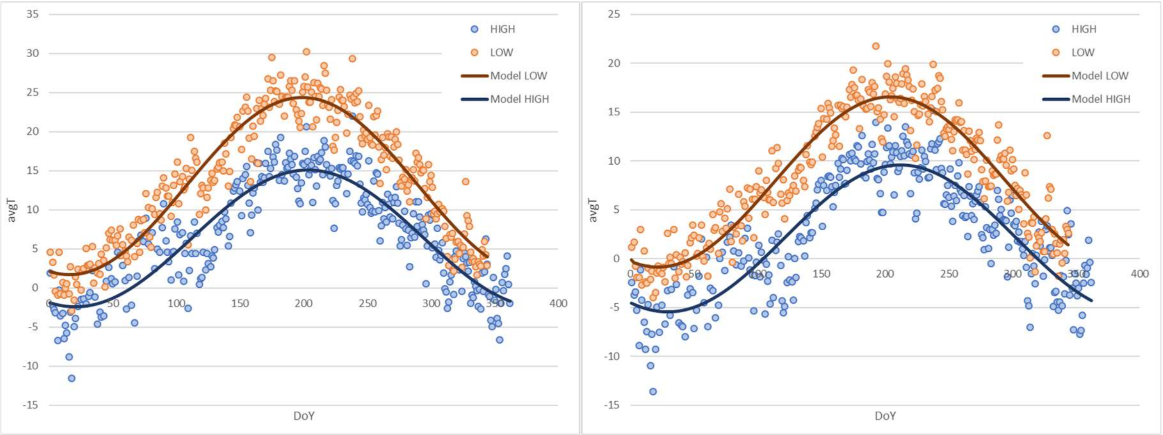


**Fig. S1.** Average day-time (left) and night-time temperatures (right), showing the measured data (points) gathered by weather stations at low (910m) and high elevations (2135m), and the model fits (lines) using the five parameter model described in eq. S27 and Table S3.

**Table S2.** Parameter values for the day-time and night-time temperature curves described in eq. 1.

| Description | Parameter | Day-time | Minimum |
|---|---|---|---|
| Average yearly temperature at 0m elevation | *avgYT* | 18.1 | 10.8 |
| Change in Average yearly temperature with a 1000m increase in elevation | *dYT* | -5.5 | -4.99 |
| Base amplitude at 0m elevation | *A0* | 31.33 | 20.0 |
| Change in amplitude with a 1000m increase in elevation | *dA* | -2.1 | -0.62 |
| Base phase at 0m elevation | *Ph0* | 260.62 | 250.1 |
| Change in phase with a 1000m increase in elevation | *dPh* | -3.61 | -2.74 |
| Base standard deviation around the mean at 0m elevation | *sdT0* | 2.9 | 2.5 |
| Change in phase with a 1000m increase in elevation | *dsdT* | 0 | 0.87 |

*Germination*

Germination (*G*, days) is simulated as being dependent on the plant genotype (GM, dimensionless), temperature (T, ºC), and two parameters that denote the maximum effect of genotype on germination (uG, days), and the effect of temperature on germination (uT, ºC days; calibrated based on the germination rates measured in the germination experiment (see Fig. S6)).

$$G = GM * uG + \frac{uT}{T} \tag{S3}$$

*Plant structure*

The cost of producing a new stalk is determined by the plant's stalk height trait ($SH_p$), which determines the diameter required to support the stalk ($D_p$, m) and the number of stalk leaves ($nSL_p$). The stalk diameter is calculated using the stalk height trait ($SH_p$), the maximum stalk height (*SHmax,* m), and two parameters derived from measured data on stem diameter and stem height in field grown plants (a=1.26, b=823.27, Fig. S7, following the relationship described in Niklas 1993).

$$D_p = \sqrt[a]{\frac{SH_p * SHmax}{b}} \quad \text{(S4)}$$

The number of stalk leaves (*$nSL_p$*) is calculated using the stalk height trait (*$SH_p$*) and a parameter that denotes the number of stalk leaves per meter stalk height (*nSL0*), which was derived from data on field grown plants.

$$nSL_p = nSL0 * SH_p * SHmax \quad \text{(S5)}$$

The stalk height, stalk diameter and the number of stalk leaves can then be combined to calculate the carbon cost of producing a stalk (*cS*, g) using a parameter that denotes the cost of producing a flower head (*cF*, g), the stalk tissue density (*STD,* g $m^{-3}$), the length of stalk leaves (*SLL*, m), the leaf width (*LW*, m), and the specific leaf area (*SLA*, $m^2$ $g^{-1}$).

$$cS_p = cF + SH_p * SHmax * \pi * \left(\frac{D_p}{2}\right)^2 * STD + nSL_p * \frac{SLL * LW}{SLA} \quad \text{(S6)}$$

The simulated plants form a rosette of long, slender leaves. In the model the length of the longest leaf (*$LLL_p$,* m) was determined by the total leaf biomass (*$Bio_{L,p}$*, g), the maximum leaf length (*LLLmax*, m), and two parameters derived from data on the relationship between rosette biomass and rosette radius (*a*=0.2774*; b*=0.0053, Fig. S8).

$$LLL_p = \min(LLLmax, a * Bio_{L,p} + b) \quad \text{(S7)}$$

We assumed that leaf length distribution within the rosette scaled linearly with leaf rank and ranged from LLL to zero. Therefore, the number of leaves in the rosette was equal to twice the number of leaves if all leaves had a length of LLL. The number of leaves in the rosette (*$nL_p$*) was thus calculated with the length of the longest leaf, the total leaf biomass, the specific leaf area (*SLA*, $m^2$ $g^{-1}$) and the leaf width (*LW*, m), which was assumed to be constant over all leaves regardless of leaf length or leaf age.

$$nL_p = 2 * Bio_{L,p} * \frac{SLA}{LLL_p * LW} \tag{S8}$$

The length of a leaf ($LLen_{l,p}$, m) was then calculated with the leaf's rank ($rank_{l,p}$, ranging from zero to $nL_p$-1), the length of the longest leaf and the total number of leaves, such that the leaf with the lowest rank was the longest leaf.

$$LLen_{l,p} = LLL_p * \frac{nL_p - rank_{l,p} + 1}{nL_p + 1} \tag{S9}$$

These leaves were growing from a central point in the rosette following a spiral phyllotaxis (*Phyl* = 137.5 degrees), at a random insertion angle ($angle_{l,p}$, degrees from the vertical) ranging from 5-85 degrees from the vertical.

*Frost damage*

Frost damage can occur when the night-time minimum temperature falls below the freezing tolerance threshold, leading to frost damage to the leaves, and potentially to plant mortality. The effects of frost on leaves and survival were expressed as the fraction of damaged leaf area, and the probability of mortality, respectively. Frost damage ($F_p$, *dimensionless; 0-1*) is calculated with the minimum night-time temperature ($T_{min}$, ºC) and two parameters; the minimum temperature at which 50% frost damage occurs (*LT50,* ºC), and the steepness of the temperature-frost damage curve (*sLT*), which determines the range from 0 to 100% frost damage.

$$F_p = \frac{1}{1 + e^{sLT*(T_{min} - LT50)}} \tag{S10}$$

Frost damage then reducing leaf biomass, leaf length and leaf nitrogen.

$$Bio_{l,p} = \left(1 - F_p\right) * Bio_{l,p} \tag{S11}$$

$$LL_{l,p} = \left(1 - F_p\right) * LL_{l,p} \tag{S12}$$

$$N_{l,p} = \left(1 - F_p\right) * N_{l,p} \tag{S13}$$

Frost damage was also used to describe the probability that the plant would die as a result of the freezing event (i.e. $F_p$ = probability of mortality), in which case the plant was removed from the simulation.

*Roots and nitrogen uptake*

The root system is modelled as a cone with a radius ($RR_p$, m), length (i.e. the rooting depth, $RD_p$, m) and volume ($RV_p$, $m^3$), which depend on the biomass of the root system ($Bio_{R,p}$, g). The relationship between biomass and root system size is parameterised using the root architectural model described in Pagès et al. (2014), using the parameterisation presented for *Pisum savium* (Table S3, Fig. S2).

$$RR_p = \frac{0.0985 * Bio_{R,p}}{(0.1 + Bio_{R,p})} + 0.032 * Bio_{R,p} + 0.0286 \tag{S14}$$

$$RD_p = \frac{0.223 * Bio_{R,p}}{(0.15 + Bio_{R,p})} + 0.015 * Bio_{R,p} \tag{S15}$$

$$RV_p = \frac{RD_p * \pi * RR_p^2}{3} \tag{S16}$$

**Table S3.** Parameter values used to derive the relationship between root system mass and root system volume, based on the *Pisum savium* model parameterisation reported in Pagès et al. (2014).

| Description | Abbreviation | Value |
|---|---|---|
| Initial root diameter | *Di* | $0.5 * 10^{-3}$ |
| Minimum root diameter | *Dmin* | $0.1 * 10^{-3}$ |
| Ratio between mother and daughter roots | *RDM* | 0.5 |
| Root tissue density | *RTD* | $0.2 * 10^{6}$ |

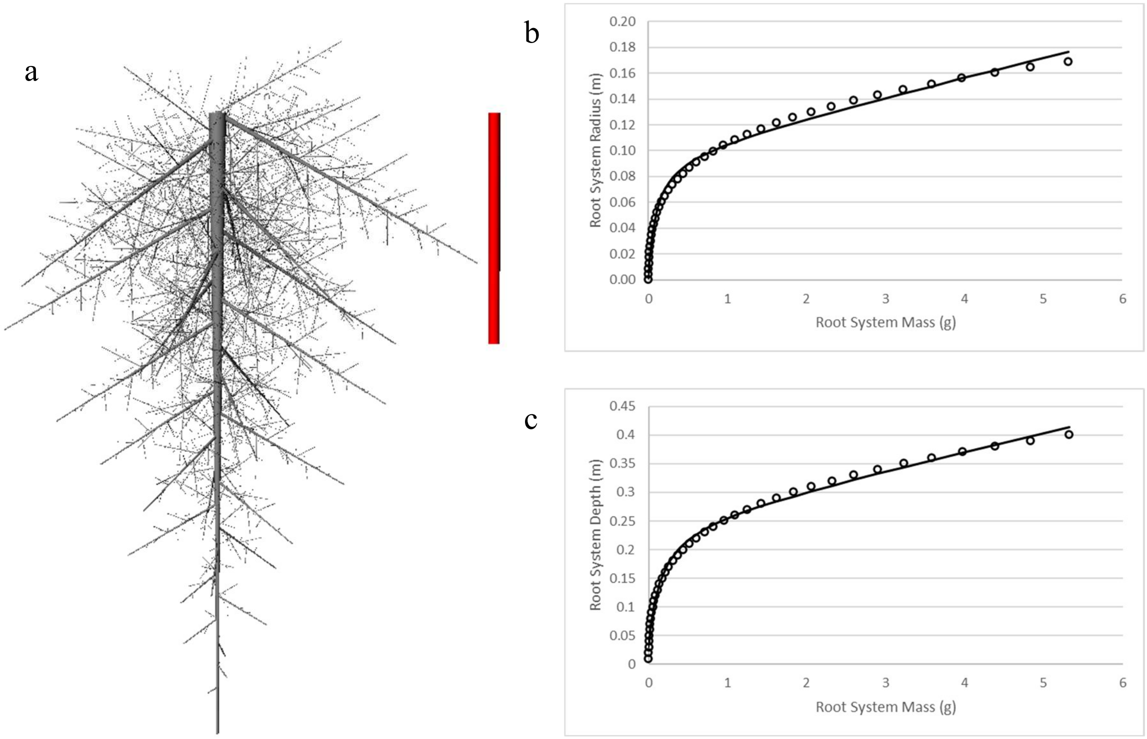


**Fig. S2.** Visualization of the root architectural model described in Pagès et al. (2014) (a, red bar = 10cm), which was used to parameterize the relationship between the mass of the root system (g, x-axes), root system depth (m, b) and root system radius (m, c). The points show the output of the root architectural model, the lines show the fitted relationship that was used in this study to describe the root system as a conical volume who's size is determine by the root system mass.

We assumed that the plant can take up only mineral nitrogen (*Nm*, g), which we assumed to be homogeneously distributed in the soil matrix due to the high mobility of mineral nitrogen. We further assumed that the mineral nitrogen content of the soil is zero at the start of the growing season and increases throughout the growing season through mineralisation by soil biota, which is modelled as a function of temperature. The mineral nitrogen content of the soil (*Nm*, g $m^{-3}$) is calculated using the model described in Stanford and Smith (1972), which requires the potentially mineralizable nitrogen content of the soil (*Nm0*, g $m^{-3}$), a mineralisation constant (*k*, $day^{-1}$), and time (*t*, days).

$$Nm = Nm0 * (1 - e^{-k*t}) \quad \text{(S17)}$$

The mineralisation constant (*k*, $day^{-1}$) is calculated using the relationship between the mineralisation constant and temperature (*T*, degrees C) reported for a forest soil with a moisture content ≥60% in Guntiñas et al. (2012) (Fig. S3).

$$k = 0.001836 * (1 - e^{0.085588*T}) \quad \text{(S18)}$$

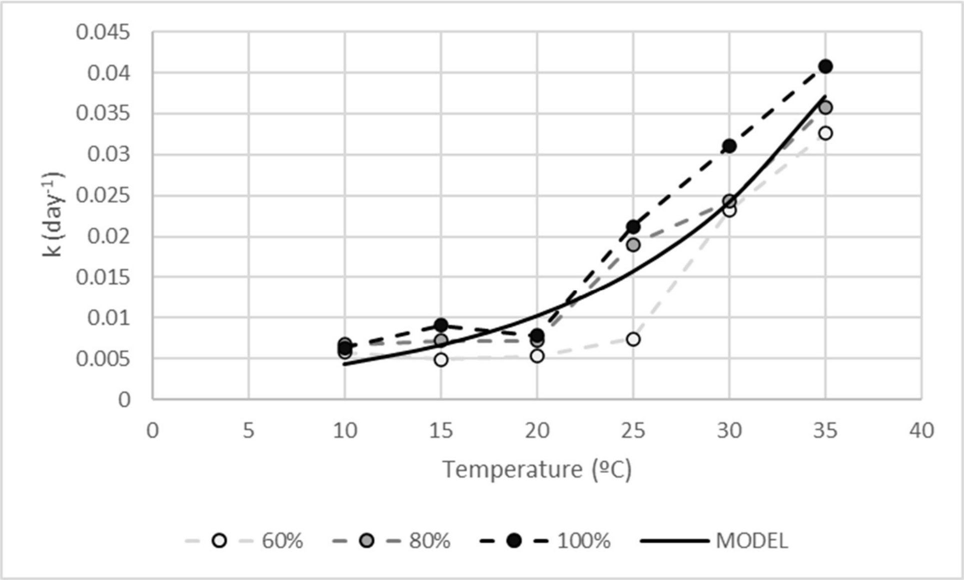


**Fig. S3.** Relationship between temperature and mineralisation constant (k, $day^{-1}$), for forest soils (dashed lines) with a moisture content of 60% (white), 80% (grey) and 100% (black) of field capacity (Guntiñas et al. 2012), and the model fit used to describe this relationship (solid line).

The potential nitrogen uptake rate ($Upot_p$, g N $day^{-1}$) of the root system can be calculated with the mineral nitrogen in the soil (*Nm*, g $m^{-3}$) and the volume of the root system ($RV_p$, $m^3$) and the time it takes for the mineral nitrogen to homogeneously distribute in the soil, which we assumed to be 1 day ($d$ = 1 day). If the total potential nitrogen uptake of all plants is higher than the soil nitrogen availability, the available nitrogen was distributed between plants based on their root volume.

$$Upot_p = \min\left(\frac{Nm * RV_p}{d}, \frac{Nm}{d} * \frac{RV_p}{\sum_{p=1}^{n} RV_p}\right) \qquad \text{(S19)}$$

*Photosynthesis*

Light interception of the leaves was calculated using the Monte-Carlo pathtracer embedded in GroImp. The light environment was simulated though both randomly arranged diffuse light sources and direct light sources spread over the solar path (Evers et al. 2010, de Vries et al. 2018) at a latitude corresponding to Switzerland. Plots of simulated plants were replicated 25 times in the x and y directions for light model calculations, creating 625 identical copies of every individual plant organ. The light conditions experienced by these 625 copies were then averaged to calculate the light interception for every individual plant organ, a procedure which effectively eliminated border effects in light interception. Leaf photosynthesis was calculated using a Farquarhar, von Caemmerer and Berry model (Farquhar et al. 1980) driven by leaf light interception and temperature. For a detailed model description, see Yin et al. (2009). In this study, we used the indicative values for constants used in the $C_3$ photosynthesis model provided in Yin and Struik (2009).

The $CO_2$ assimilated by the leaves ($A_{l,p}$, µmol $CO_2$ m$^{-2}$ s$^{-1}$) was converted to biomass ($Bio_{A,p}$, g d$^{-1}$) using leaf length ($LL_{l,p}$, m), leaf width ($LW$, m), a conversion parameter ($C$) to convert from µmol $CO_2$ to grams of glucose and seconds to days, and a second conversion parameter accounted for the construction costs from glucose to biomass production ($cc$, g biomass g$^{-1}$ glucose).

$$Bio_{A,p} = C * cc * \sum_{l=1}^{n} (A_{l,p} * LL_{l,p} * LW) \tag{S20}$$

*Plant growth*

We assumed that the C:N ratio of plant tissues is conserved irrespective of the availability of carbon and nitrogen, This assumption allows us to describe the amount of biomass available for growth ($Bio_{G,p}$, g d$^{-1}$) as a function that is limited by either the potential nitrogen uptake of the root system ($Upot_p$, g d$^{-1}$) and the assumed nitrogen content of plant biomass (*Nbio,* g g$^{-1}$), or the assimilates produced by photosynthesis ($Bio_{A,p}$, g d$^{-1}$), minus the costs of maintenance respiration, which is a function of the maintenance respiration rate (*rm*, g C g$^{-1}$ N d$^{-1}$) and the total plant nitrogen content ($N_p$, g).

$$Bio_{G,p} = \min\left(\frac{Upot_p}{Nbio}, Bio_{A,p} - rm * N_p\right) \tag{S21}$$

The realised nitrogen uptake rate ($U_p$, g d$^{-1}$) was then calculated based on plant growth ($Bio_{G,p}$, g d$^{-1}$).

$$U_p = Bio_{G,p} * Nbio \tag{S22}$$

The realised nitrogen uptake rate ($U_p$, g d$^{-1}$) was then multiplied by the time step (*dt,* d) and added to plant's nitrogen pool ($N_p$, g).

$$N_p = N_p + U_p * dt \tag{S23}$$

The allocation of biomass to plant growth is dependent on plant phenology. The plant is considered to be in its vegetative stage of development if the growing degree days accumulated by the plant since germination ($sumGDD_p$, gdd) do not exceed the flowering threshold set by the time to flowering trait ($FL_p$, gdd) and the maximum time to flowering (*FLmax*, gdd). In this vegetative stage ($cumGDD_p<FL_i*FLmax$), biomass allocated to growth is distributed between

root ($Bio_{R,p}$, g) and leaf biomass ($Bio_{L,p}$, g), assuming a conserved root:leaf ratio ($RLR$, g g$^{-1}$).

$$\frac{dBio_{R,p}}{dt} = Bio_{G,p} * \frac{RLR}{RLR+1} \qquad if\left(sumGDD_p < FL_p * FLmax\right)$$

(S24)

$$\frac{dBio_{L,p}}{dt} = Bio_{G,p} * \left(1 - \frac{RLR}{RLR+1}\right) \qquad if\left(sumGDD_p < FL_p * FLmax\right)$$

In the generative stage of development (i.e. when $sumGDD_p \geq FL_i * FLmax$), biomass allocated to growth is distributed between stalk ($Bio_{ST,p}$, g) and seed biomass ($Bio_{SE,p}$, g) following a hierarchical allocation model that prioritizes allocation to seeds over allocation to stalks. Allocation to seeds and stalks is calculated using the seed sink strength ($Sink_{SE,p}$, g), which was defined as the difference between the current seed biomass ($Bio_{SE,p}$, g) and the total plant seed set ($SE_p$, g).

$$Sink_{SE,p} = SE_p - Bio_{SE,p} \qquad \text{(S25)}$$

$$\frac{dBio_{ST,p}}{dt} = Bio_{G,p} * \frac{Sink_{ST,p}}{Sink_{ST,p} + Sink_{SE,p}} \qquad if\left(cumGDD_p \geq FL_p * FLmax\right) \quad \text{(S26)}$$

$$\frac{dBio_{SE,p}}{dt} = Bio_{G,p} * \frac{Sink_{SE,p}}{Sink_{ST,p} + Sink_{SE,p}} \qquad if\left(cumGDD_p \geq FL_p * FLmax\right)$$

*Pollination*

To simulate pollinator preference for more apparent flowers, we used the light intercepted by the flower as a proxy for flower attractiveness, so that flowers at the top of the canopy are more attractive than flowers lower in the canopy. The pollinator visits paid to a flower ($P_f$, number of visits) was thus calculated with the pollinator abundance ($P$, number of visits m$^{-2}$ h$^{-1}$), the surface area of the plot (area, m$^2$), the daytime hours in this time step ($h$, h), the light interception by the flower ($APAR_f$, µmol m$^{-2}$ s$^{-1}$), and the total attractiveness of all flowers in the plot.

$$P_f = P * area * h * \frac{APAR_f}{\sum_{f=1}^{n} APAR_f} \qquad \text{(S27)}$$

Seed set was shown to be pollen limited and followed a saturating dose-response curve function with a threshold of 50 deposited pollen grains to initiate seed development (Bloch et al. 2006). In the model, total plant seed set ($SE_{,p}$, g) was calculated with the pollinator visits of each inflorescence ($P_{s,p}$, visits), the pollen deposition rate ($p$, pollen visit$^{-1}$), the maximum number of seeds per inflorescence ($smax$, seeds inflorescence $^{-1}$), the seed weight ($sw$, g), and two parameters that were derived from Bloch et al. (2006) (*a=14.752, b=58.035*, Fig. S4).

$$SE_p = \sum_{s=1}^{n} \left(smax * \left(a * \log\left(P_{f,p} * p\right) - b\right) * sw\right) \qquad \text{(S28)}$$

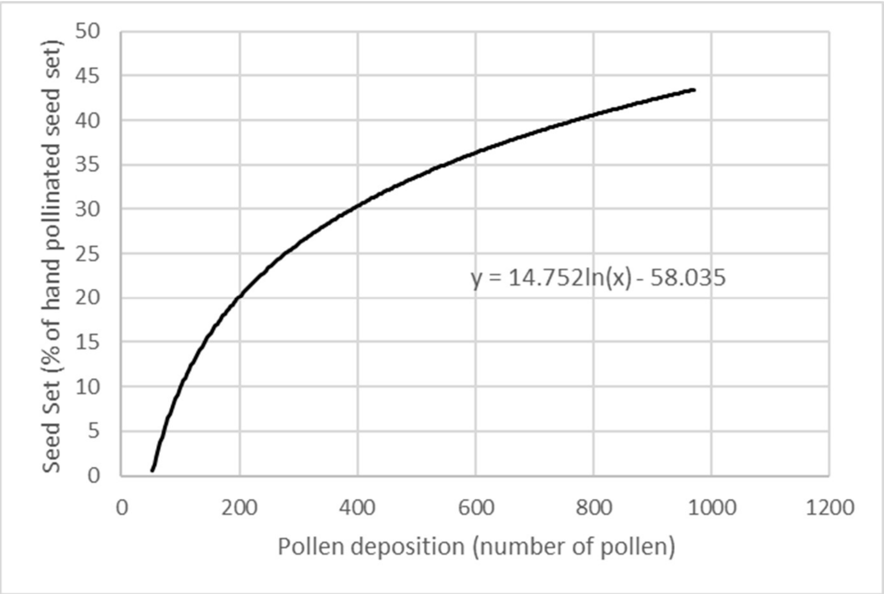


**Fig. S4.** Relationship between pollen deposition and seed set as a % of the hand pollinated seed set based on the results presented in Bloch et al. (2006).

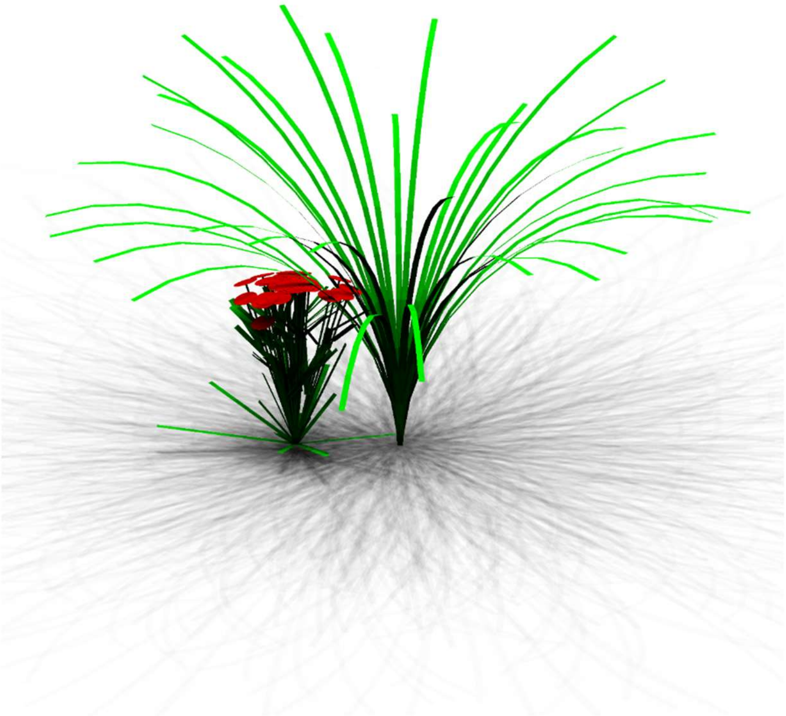

**Fig. S5.** Visualisation of the tall grass architecture used in the model, compared to a plant from the high elevation habitat.

**Table S4.** List of model parameters and their values.

| Description | Name | Value | Unit (reference) | Eq. |
|---|---|---|---|---|
| Root leaf ratio | RLR | 1 | g $g^{-1}$ | S2 |
| Maximum effect of genotype on germination | uG | 20 | days | S3 |
| Effect of temperature on germination | uT | -70.335 | ºC days | S3 |
| Maximum stalk length | SLmax | 1[a] | m | S4-6,21 |
| Stalk tissue density | STD | 250000[b] | g $m^{-3}$ | S4-6,21 |
| Maximum leaf length | LLLmax | 0.15[a] | m | S7 |
| Specific leaf area | SLA | 0.01459[c] | $m^2$ $g^{-1}$ | S8 |
| Leaf width | LW | 0.005[a] | m | S8 |
| Minimum temperature at which 50% frost damage occurred | LT50 | -8 | ºC | S10 |
| Steepness of the temperature – frost damage curve | sLT | 1 | ºC | S10 |
| Potentially mineralizable nitrogen content of the soil | Nm0 | 0.004[d] | g $m^{-3}$ | S17 |
| Construction costs of converting glucose to biomass | cc | 0.667[e] | g biomass $g^{-1}$ glucose | S20 |
| Maintenance respiration rate | rm | 0.218[f] | g C g $N^{-1}$ $d^{-1}$ | S21 |
| Nitrogen content of plant biomass | Nbio | 0.03 | g nitrogen $g^{-1}$ biomass | S21,22 |
| Pollen deposition rate | P | 100[g] | pollen $visit^{-1}$ | S28 |
| Maximum number of seeds per inflorescence | smax | 100[a] | seeds $inflorescence^{-1}$ | S28 |
| Seed weight | sw | 5e-4[a] | g | S28 |
| Phyllotaxis | Phyl | 137.5 | degrees | |
| Standard deviation of offspring trait value around parental trait value | Tsd | 0.05 (generation 1-75)<br>0.005 (generation 76-125) | dimensionless | |

[a] derived from data on field grown plants published in Palsson (2021); [b] representative value for forbs derived from Ishida et al. (2008); [c] Kleyer et al. (2008) ; [d] Körner (2003); [e] De Vries et al. (1974); [f] Ryan (1991) ; [g] Bloch et al. (2006);

**Results S1**

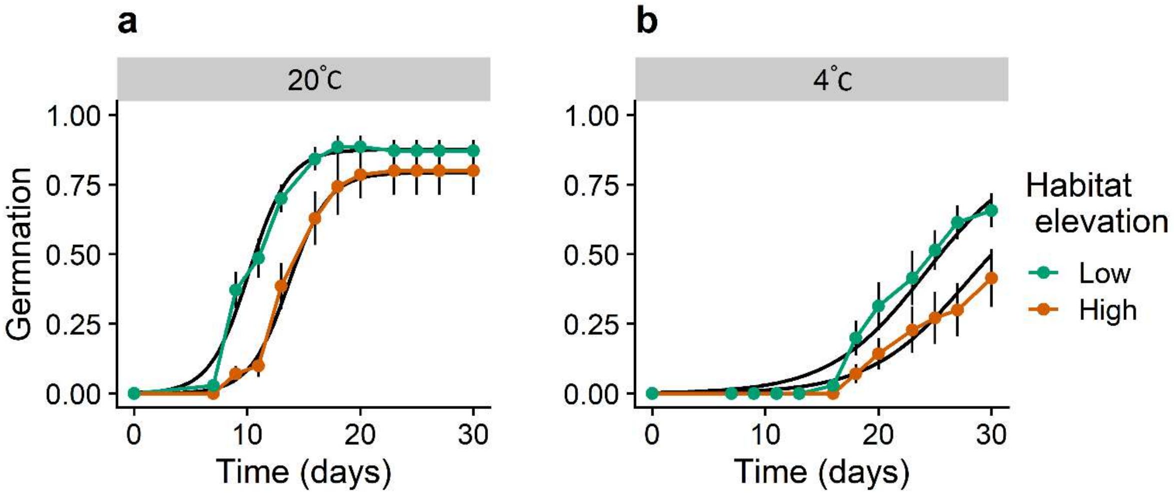


**Fig. S6.** *In vivo* germination of plants from the low (green) and high (red) elevation habitats under 20 ºC (**a**) and 4 ºC (**b**), and the model fit used in the calibration of the E-FSP model (black). Error bars show the standard error of the mean.

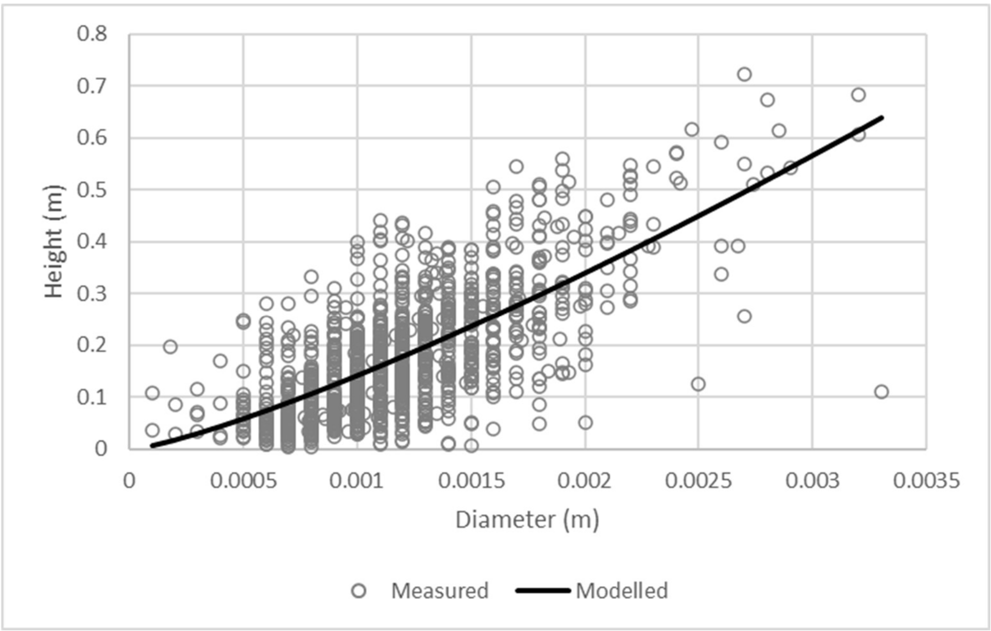


**Fig. S7.** Stalk diameter height relationship of field grown plants (points) and the relationship used in the model (line).

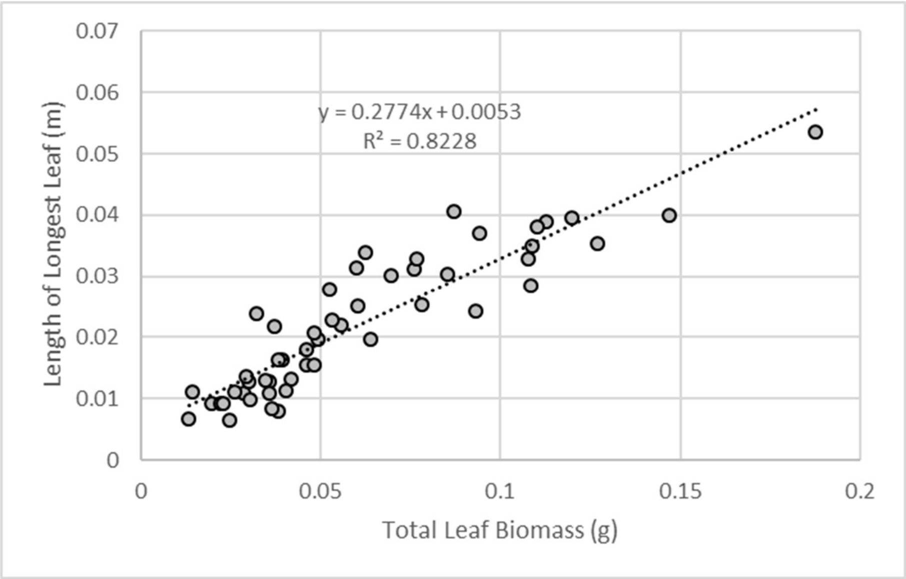


**Fig. S8.** Relationship between total leaf biomass and the length of the longest leaf in field grown plants (points), and the relationship used in the model (line).

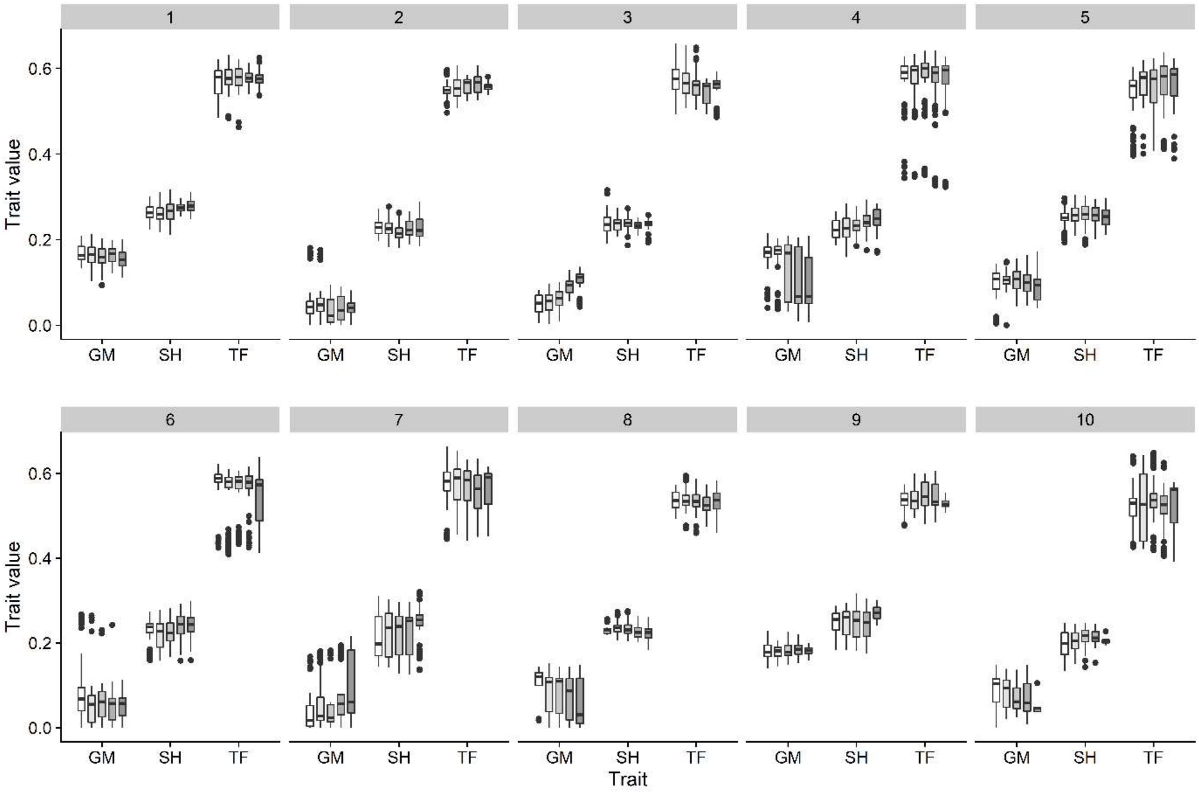


**Fig. S9.** Trait variation in the ten replicate populations of the control treatment after 105, 110, 115, 120, 125

(white to grey, respectively) (y-axis: trait value (0-1); x-axis; Germination (GM); Stalk Height (SH) and Time to Flowering (TF)).

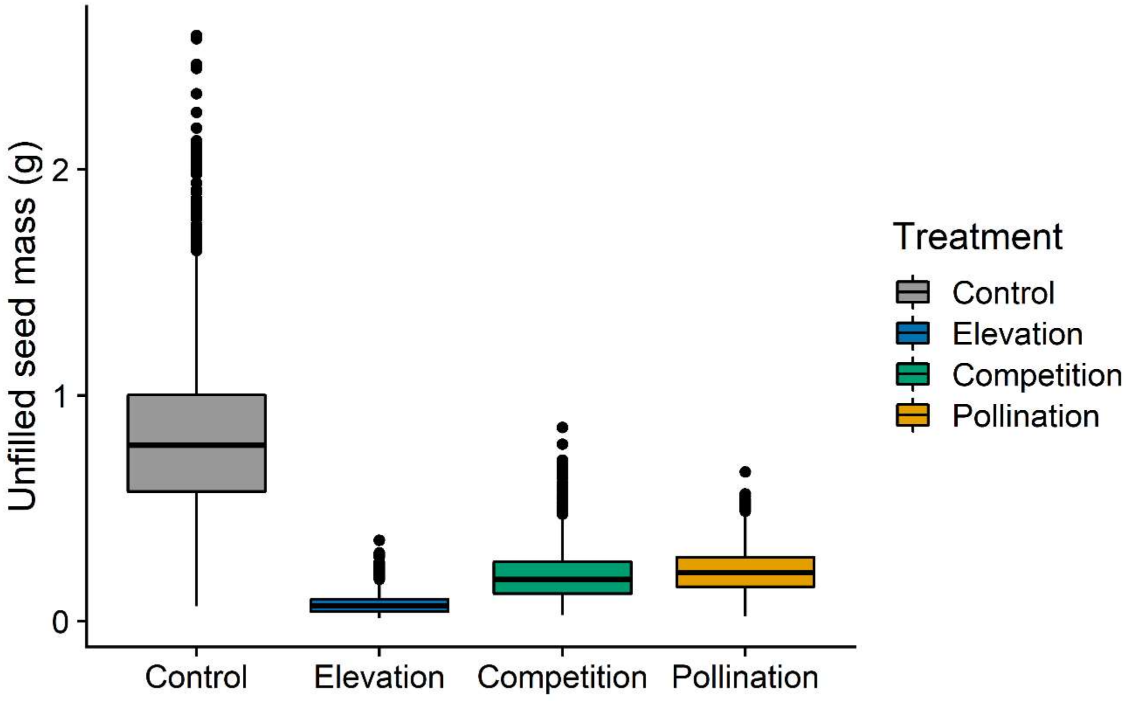


**Fig. S10.** Unfilled seed mass (g, y-axis) of individual plants in four treatments; Control, grey; Elevation, blue; Competition, green; and Pollination, yellow.

**Table S5.** Results of traits and variables of in vivo and in silico populations, expressed as mean ± standard deviation.

| Method | Trait / Variable | Low elevation | High elevation | Unit | P value |
|---|---|---|---|---|---|
| *in vivo* | Germination (GM) | 0.404±0.196 | 0.539±0.225 | *dimensionless* | <0.001 |
| | Stalk height (SH) | 0.265±0.141 | 0.087±0.059 | *dimensionless* | <0.001 |
| | Time to flowering (TF) | 0.512±0.033 | 0.416±0.025 | *dimensionless* | <0.001 |
| | Flowering time | 187±12.2 | 201.6±15.3 | Julien day | <0.001 |
| | Number of stalks | 2.45±2.47 | 1.5±2.06 | stalks | <0.001 |
| | Rosette area | 42.52±56.67 | 9.83±1.22 | $cm^2$ | <0.001 |
| | Fitness | 0.046±0.086 | 0.024±0.022 | g | <0.05 |
| *in silico* | Germination (GM) | 0.038±0.033 | 0.406±0.245 | *dimensionless* | |
| | Stalk height (SH) | 0.207±0.031 | 0.074±0.032 | *dimensionless* | |
| | Time to flowering (TF) | 0.491±0.077 | 0.282±0.039 | *dimensionless* | |
| | Flowering time | 174.6±12.1 | 222.3±9 | Julien day | |
| | Number of stalks | 4.64±3.13 | 2.99±2.6 | stalks | |
| | Rosette area | 215.8±193.5 | 9.27±6.98 | $cm^2$ | |
| | Fitness | 0.171±0.12 | 0.033±0.024 | g | |

### References


Bloch, D., N. Werdenberg, and A. Erhardt. 2006. Pollination crisis in the butterfly-pollinated wild carnation Dianthus carthusianorum? New Phytologist **169**:699-706.

De Vries, F. P., A. Brunsting, and H. Van Laar. 1974. Products, requirements and efficiency of biosynthesis a quantitative approach. Journal of Theoretical Biology **45**:339-377.

de Vries, J., J. B. Evers, E. H. Poelman, and N. P. Anten. 2020. Simulation of optimal defence against herbivores under resource limitation and competition using an evolutionary functional-structural plant model. in silico Plants **2**.

de Vries, J., E. H. Poelman, N. P. Anten, and J. B. Evers. 2018. Elucidating the interaction between light competition and herbivore feeding patterns using functional–structural plant modelling. Annals of Botany **121**:1019-1031.

Evers, J., J. Vos, X. Yin, P. Romero, P. Van Der Putten, and P. Struik. 2010. Simulation of wheat growth and development based on organ-level photosynthesis and assimilate allocation. Journal of Experimental Botany **61**:2203-2216.

Farquhar, G. D., S. v. von Caemmerer, and J. Berry. 1980. A biochemical model of photosynthetic CO 2 assimilation in leaves of C 3 species. Planta **149**:78-90.

Guntiñas, M. E., M. Leirós, C. Trasar-Cepeda, and F. Gil-Sotres. 2012. Effects of moisture and temperature on net soil nitrogen mineralization: A laboratory study. European Journal of Soil Biology **48**:73-80.

Hemmerling, R., O. Kniemeyer, D. Lanwert, W. Kurth, and G. Buck-Sorlin. 2008. The rule-based language XL and the modelling environment GroIMP illustrated with simulated tree competition. Functional Plant Biology **35**:739-750.

Ishida, A., T. Nakano, K. Yazaki, S. Matsuki, N. Koike, D. L. Lauenstein, M. Shimizu, and N. Yamashita. 2008. Coordination between leaf and stem traits related to leaf carbon gain and hydraulics across 32 drought-tolerant angiosperms. Oecologia **156**:193.

Kleyer, M., R. Bekker, I. Knevel, J. Bakker, K. Thompson, M. Sonnenschein, P. Poschlod, J. Van Groenendael, L. Klimeš, and J. Klimešová. 2008. The LEDA Traitbase: a database of life-history traits of the Northwest European flora. Journal of Ecology **96**:1266-1274.

Körner, C. 2003. Alpine plant life: functional plant ecology of high mountain ecosystems. Springer.

Niklas, K. J. 1993. Influence of Tissue Density-specific Mechanical Properties on the Scaling of Plant Height. Annals of Botany **72**:173-179.

Pagès, L., C. Bécel, H. Boukcim, D. Moreau, C. Nguyen, and A.-S. Voisin. 2014. Calibration and evaluation of ArchiSimple, a simple model of root system architecture. Ecological Modelling:76-84.

Ryan, M. G. 1991. Effects of climate change on plant respiration. Ecological applications **1**:157-167.

Stanford, G., and S. Smith. 1972. Nitrogen mineralization potentials of soils. Soil Science Society of America Journal **36**:465-472.

Yin, X., and P. Struik. 2009. C3 and C4 photosynthesis models: an overview from the perspective of crop modelling. NJAS-Wageningen Journal of Life Sciences **57**:27-38.

Yin, X., P. C. Struik, P. Romero, J. Harbinson, J. B. Evers, P. E. Van Der Putten, and J. Vos. 2009. Using combined measurements of gas exchange and chlorophyll fluorescence to estimate parameters of a biochemical C3 photosynthesis model: a critical appraisal and a new integrated approach applied to leaves in a wheat (Triticum aestivum) canopy. Plant, Cell & Environment **32**:448-464.
